# Macrophages undergo functionally significant reprogramming of nucleotide metabolism upon classical activation

**DOI:** 10.1101/2023.12.27.573447

**Authors:** Steven V. John, Gretchen L. Seim, Billy J. Erazo-Flores, John Steill, Jack Freeman, James A. Votava, Nicholas L. Arp, Xin Qing, Ron Stewart, Laura J. Knoll, Jing Fan

**Affiliations:** Morgridge Institute for Research, Madison, WI; Cellular and Molecular Biology Graduate Program, University of Wisconsin–Madison, Madison, WI; Department of Nutritional Sciences, University of Wisconsin–Madison, Madison, WI; Cellular and Molecular Pathology Graduate Program, University of Wisconsin–Madison, Madison, WI; Department of Medical Microbiology and Immunology, University of Wisconsin–Madison, Madison, WI

## Abstract

During an immune response, macrophages specifically rewire their metabolism to support functional changes. Using a multi-omics approach, we identified nucleotide metabolism as one of the most significantly rewired pathways across the metabolic network in classically activated macrophages. Further isotopic tracing studies revealed the substantial changes in nucleotide *de novo* synthesis, degradation, and salvage fluxes in stimulated macrophages, as well as the key reactions where metabolic regulation occurs: 1) *de novo* synthesis of purine nucleotides is shut down and particularly blocked at the last step of IMP synthesis catalyzed by ATIC; 2) *de novo* synthesis of pyrimidines is maintained up to UMP, but further synthesis of CTP (catalyzed by CTPS) and dTMP (catalyzed by TYMS) is greatly reduced; 3) Nucleotide degradation to nitrogenous bases is increased, but further oxidation of purine bases (catalyzed by XOR) is inhibited, causing a great accumulation of nucleosides and bases; and 4) cells switch to salvaging the nucleosides and bases as the primary means to maintain purine nucleotides. Mechanistically, we found these changes are driven by a combination of transcriptional regulation and enzyme inhibition. Nitric oxide (NO) was identified as a major regulator, driving the strong inhibition of ATIC and XOR, and the transcriptional downregulation of *Tyms*. To understand the functional impact of the activation-induced switch from purine *de novo* synthesis to salvage, we knocked out the purine salvage enzyme *Hprt*. *Hprt* knockout significantly alters functional gene expression in activated macrophages, suppresses macrophage migration, and increases pyroptosis. Furthermore, knocking out *Hprt* or *Xor* increases the proliferation of the intracellular parasite *Toxoplasma gondii* in macrophages. Together, these results comprehensively uncovered the dynamic rewiring of nucleotide metabolism in classically activated macrophages, elucidated the key regulatory mechanisms, and identified the functional significance of such rewiring.

## Introduction

Macrophages are crucial cells of the innate immune system with great functional plasticity. Macrophages can both take on a variety of functional states in response to different molecular signals associated with diverse biological scenarios (e.g., infection, injury, and inflammation) and dynamically change their functions over the course of an immune response. For instance, upon sensing classical activation signals associated with infection such as lipopolysaccharide (LPS, a bacterial cell wall component) and interferon-γ (IFNγ), macrophages first turn on functions that are pro-inflammatory to help eliminate pathogens. Then over time, these responses subside, and macrophages develop more anti-inflammatory responses to help promote healing (Foster and Medzhitov, 2009; Ivashkiv, 2011; Hotchkiss et al., 2013; Murray, 2017).

Dynamic reprogramming of metabolism plays a critical and increasingly recognized role in the functional transitions in macrophages during an immune response. Over the past decade, growing efforts have sought to identify the *metabolic characteristics* specifically associated with different functional states of macrophages and to elucidate the *metabolic tipping points* where alteration of key metabolic activities can orchestrate macrophage functions. Such efforts have proven fruitful and have broad significance, especially given macrophages’ wide-ranging involvement in many health conditions. One of the most demonstrated examples is the citrate cycle: differential and dynamic rewiring of multiple steps in the citrate cycle were found to be specifically associated with different functional states in macrophages, and such rewiring is important for both supporting and regulating macrophage immune responses (reviewed in Ryan and O’Neill, 2020; He et al., 2021; Seim and Fan, 2022). Increasing evidence suggests that the metabolic rewiring associated with macrophage immune activation is system-wide, occurring throughout metabolic networks (Hseih et al., 2020). However, beyond central carbohydrate and lipid metabolic pathways, where many exciting discoveries have been made in macrophages over the recent years, further insights across metabolic networks remain to be elucidated.

Comprehensive understanding of metabolic reprogramming in macrophages requires answering three major questions: (1) *What* metabolic pathways or activities are substantially altered during a macrophage immune response (i.e., the metabolic characteristics)? (2) *How* do these metabolic transitions occur (i.e., the regulatory mechanism)? And (3) *Why* does such metabolic rewiring happen during an immune response, and how does it affect macrophage functions (i.e., the functional significance)? Here, we used unbiased multi-omics analysis to characterize the dynamic metabolic reprogramming in macrophages upon classical LPS+IFNγ stimulation, which identified nucleotide metabolism, a fundamental yet largely understudied pathway in the system of macrophages, as one of the most substantially remodeled metabolic pathways. Then through a series of extensive isotopic tracing studies, we revealed the profound changes in metabolic fluxes through purine and pyrimidine *de novo* synthesis, degradation, and salvage pathways, and identified key regulation points (i.e., key reactions and enzymes) whose alterations drive the systemic rewiring of nucleotide metabolism. We further elucidated the mechanisms regulating these key reactions. Finally, we demonstrated that such rewiring of nucleotide metabolism (specifically the switch from purine *de novo* synthesis to salvage upon activation) is important for various macrophage functions, including migration, pyroptosis, and interactions with intracellular parasites. Together, this study thoroughly elucidates the what, the how, and the why of nucleotide metabolism rewiring during the macrophage immune response. The discoveries here have broad significance both in relevance to health conditions closely involving macrophage responses, such as infection and inflammation, and in relevance to the novel biochemical knowledge regarding the regulation of nucleotide metabolism, which is ubiquitously important across cell types.

## Results

### Systematic reprogramming of nucleotide metabolism in macrophages upon classical activation

#### Multi-omics analysis identifies nucleotide metabolism as a substantially rewired metabolic pathway upon classical stimulation

To systematically identify the metabolic pathways that are rewired in macrophages upon classical activation, we first performed metabolomic analysis comparing murine bone marrow-derived macrophages (BMDMs) stimulated with LPS+IFNγ for 24h to unstimulated BMDMs. Pathway enrichment analysis (Xia et al., 2009) highlighted the most significantly and profoundly impacted pathways, which include several groups (Figure 1A): (1) arginine metabolism pathway, whose rewiring is one of the most well-recognized markers of macrophage polarization (Rath et al., 2014); (2) citrate cycle, which has been highlighted by recent studies for its key role in macrophage immune responses (Tannahill et al., 2013; Jha et al., 2015; Seim et al., 2019; Ryan and O’Neill, 2020); (3) purine metabolism, pyrimidine metabolism, and the pentose phosphate pathway (PPP), which are all parts of nucleotide metabolism; and (4) aspartate, alanine, and glutamine metabolism, which are closely connected to both the citrate cycle and nucleotide metabolism (Figure 1B). Overall, this analysis not only is highly consistent with prior knowledge, but also identified nucleotide metabolism, which has not been thoroughly studied in macrophages, as a crucial metabolic pathway whose substantial rewiring is strongly associated with classical activation.

**Figure 1:**
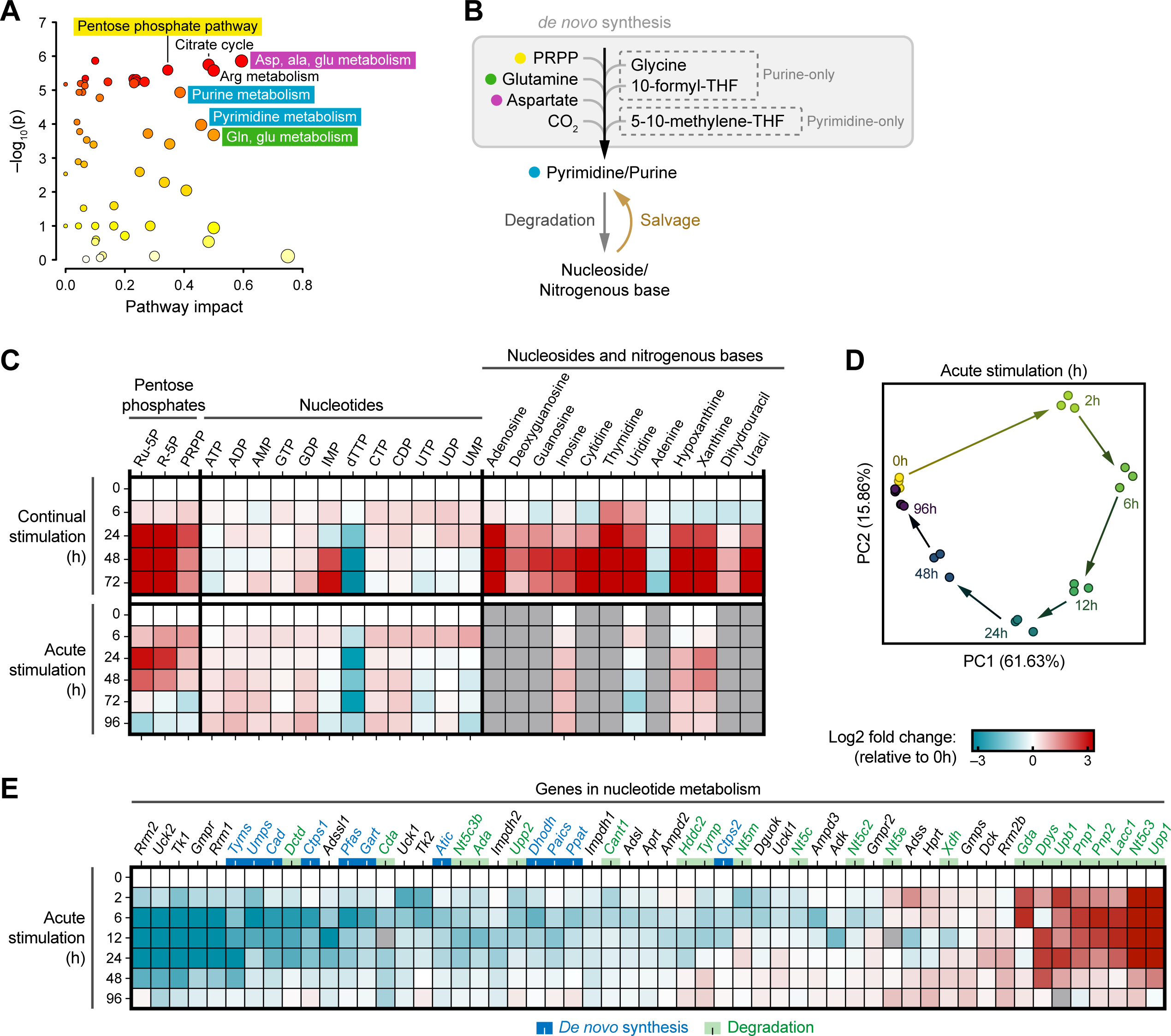
Multi-omics analysis identifies nucleotide metabolism as a substantially rewired metabolic pathway upon classical stimulation. A) Pathway enrichment analysis comparing metabolomic changes in unstimulated and 24h continually stimulated BMDMs. B) Several pathways identified in Figure 1A are directly connected to the nucleotide metabolism process. C) Metabolomic changes in pentose phosphates, nucleotides, nucleosides, and nitrogenous bases in BMDMs over a timecourse of continual or acute stimulation. D) Principal component analysis of transcriptomic changes in BMDMs over a timecourse of acute stimulation. E) Transcriptional changes in nucleotide metabolism in BMDMs over a timecourse of acute stimulation. Genes involved in nucleotide *de novo* synthesis (blue) or degradation (green) are indicated by color codes of gene names. C, E) Relative (C) metabolite abundances or (E) gene expression are compared to unstimulated cells (0h) and displayed on a log2 scale as a heatmap. Each box represents the mean of n=3 independent samples.

Given the growing appreciation for the highly dynamic nature of metabolic rewiring during macrophages’ immune response (Luan and Horng, 2021; Seim and Fan, 2022), we further profiled the changes in nucleotide metabolism over a timecourse upon either continual or acute stimulation with classical activation signals. (For continual stimulation, macrophages were constantly exposed to LPS+IFNγ over several days. For acute stimulation, macrophages were stimulated with LPS+IFNγ for only 2h, then the stimulants were removed, and cells were cultured in standard conditions over time.) Dynamic changes were observed in compounds involved in nucleotide metabolism, including pentose phosphates, nucleotides, nucleosides, and nitrogenous bases. Pentose phosphates accumulated substantially upon stimulation. Such accumulation is reversible upon acute stimulation (peaking at 24h) but was maintained upon continual stimulation (Figure 1C). The majority of nucleotides remained largely steady upon both continual and acute stimulation, with two notable exceptions: (1) Inosine monophosphate (IMP), the hub for both *de novo* synthesis and salvage of purine nucleotides, built up 10-fold after 48–72h of continual stimulation; (2) dTTP depleted over time to undetectable levels after 48h upon continual stimulation, and transiently depleted around 24–72h after acute stimulation, but began to recover at 96h (Figure 1C). Most nucleosides and nitrogenous bases, which are associated with nucleotide degradation and salvage, increased substantially and remained highly elevated (up to 10-fold or more) after 24h continual stimulation (Figure 1C). Upon acute stimulation, accumulation was also observed in inosine, hypoxanthine, and xanthine, peaking at 24–48h post-stimulation (Figure 1C). (Many other nucleosides were not well detected upon acute stimulation due to their low baseline abundance.) Overall, the metabolomics analysis revealed that both continual and acute stimulation induce substantial rewiring of macrophage nucleotide metabolism after 24h, while the changes are generally stronger when the cells are continually exposed to the stimuli. The changes upon acute stimulation indicate that the rewiring of nucleotide metabolism is largely reversible after the stimuli are removed for a long period (72–96h).

Interestingly, this substantial metabolomic transition after 24h showed temporal correlation with changes in functional markers, such as the production of cytokines. Upon continual stimulation with LPS+IFNγ, macrophages initially produce large amounts of pro-inflammatory cytokines at early time points, but at later time points (around 24–48h), such pro-inflammatory cytokine production subsides, and cells instead produce anti-inflammatory cytokines such as IL-10 (Supplemental figure 1A) (Seim et al., 2019). The fact that nucleotide metabolism rewiring occurs during this functional transition points to its potential importance. Substantial rewiring of nucleotide metabolism is commonly associated with changes in cell cycle. However, it is worth noting that BMDMs, whether stimulated or not, do not proliferate and remain highly viable over the timecourse of both continual and acute stimulation (Supplemental figures 1B–C). Interestingly, we also observed similar changes in nucleotide metabolism compounds upon continual LPS+IFNγ stimulation in another widely used macrophage cell model, the RAW 264.7 cell line. This includes the depletion of dTTP, and the strong accumulation of IMP, pentose phosphates, nucleosides, and nitrogenous bases (Supplemental figure 1D), which all temporally correlated with the increased production of anti-inflammatory cytokines. However, unlike the terminally differentiated BMDMs, RAW 264.7 cells do proliferate in the unstimulated state but stop proliferation upon stimulation (Supplemental figure 1E). These results demonstrate that the dynamic rewiring of nucleotide metabolism is consistently associated with the macrophage response to classical stimulation across cell models. Although such nucleotide metabolism rewiring is not due to changes in proliferation state, it may cause growth arrest, as observed in RAW 264.7 cells.

To comprehensively examine the dynamic changes in macrophage metabolism and function upon classical stimulation, we further conducted transcriptomic analysis in BMDMs over a timecourse upon acute stimulation. Principal component analysis reveals the gene expression profile evolved nicely over the timecourse and returned to near baseline at 96h post-stimulation (Figure 1D). Among the top 10 genes most aligned with the major dynamic changes represented by principal component 1 (PC1) are many immune response genes (*Cxcl10, Il12b, Cd40* etc.). Also included are important metabolic genes such as *Nos2* (the well-recognized metabolic marker for classical activation which functions in arginine metabolism), *Irg1* (an enzyme branching out of the citrate cycle that has been the focus of many recent studies for its importance in immunometabolism), and the nucleotide degradation gene *Upp1* (uridine phosphorylase) (Supplemental figure 1F). This highlights the temporal correlation between functional changes and metabolic rewiring across the immune response timecourse. Several nucleotide degradation genes, such as *Pnp1* and *Pnp2* (purine nucleoside phosphorylase) showed similar dynamic trends as *Upp1*—a large increase in expression upon acute stimulation that returned to baseline over time. Conversely, many genes in nucleotide *de novo* synthesis showed the opposite trend: a great reduction 6–48h post-stimulation that returned to baseline by 96h (Figure 1E). The GO term “regulation of nucleobase-containing compound metabolic process” is one of the highly enriched GO terms for its dynamic changes along the first principal component over the timecourse (adjusted p=1.47 x 10^-9^). Taken together, the transcriptomic and metabolomic results are highly consistent and show that nucleotide metabolism undergoes substantial reprogramming upon classical stimulation.

#### Nucleotide *de novo* synthesis is shut off upon stimulation

Building on the multi-omics analysis, we next sought to understand how metabolic fluxes through various nucleotide metabolism pathways are altered, which may drive the observed accumulation or depletion of metabolites upon stimulation. First, to examine nucleotide *de novo* synthesis flux, we performed kinetic isotopic tracing on stimulated BMDMs (continually stimulated with LPS+IFNγ for 48h) or unstimulated BMDMs (cultured for the same duration) using a γ-^15^N-glutamine tracer, because glutamine is a main nitrogen donor for both purine and pyrimidine *de novo* synthesis (Figure 2A). Within 2h after switching cells to media containing labeled glutamine, intracellular glutamine becomes fully labeled in both stimulated and unstimulated macrophages (Figure 2B). Likewise, the pyrimidine synthesis intermediates carbamoyl aspartate and orotate quickly become nearly fully labeled (Figure 2C).

**Figure 2:**
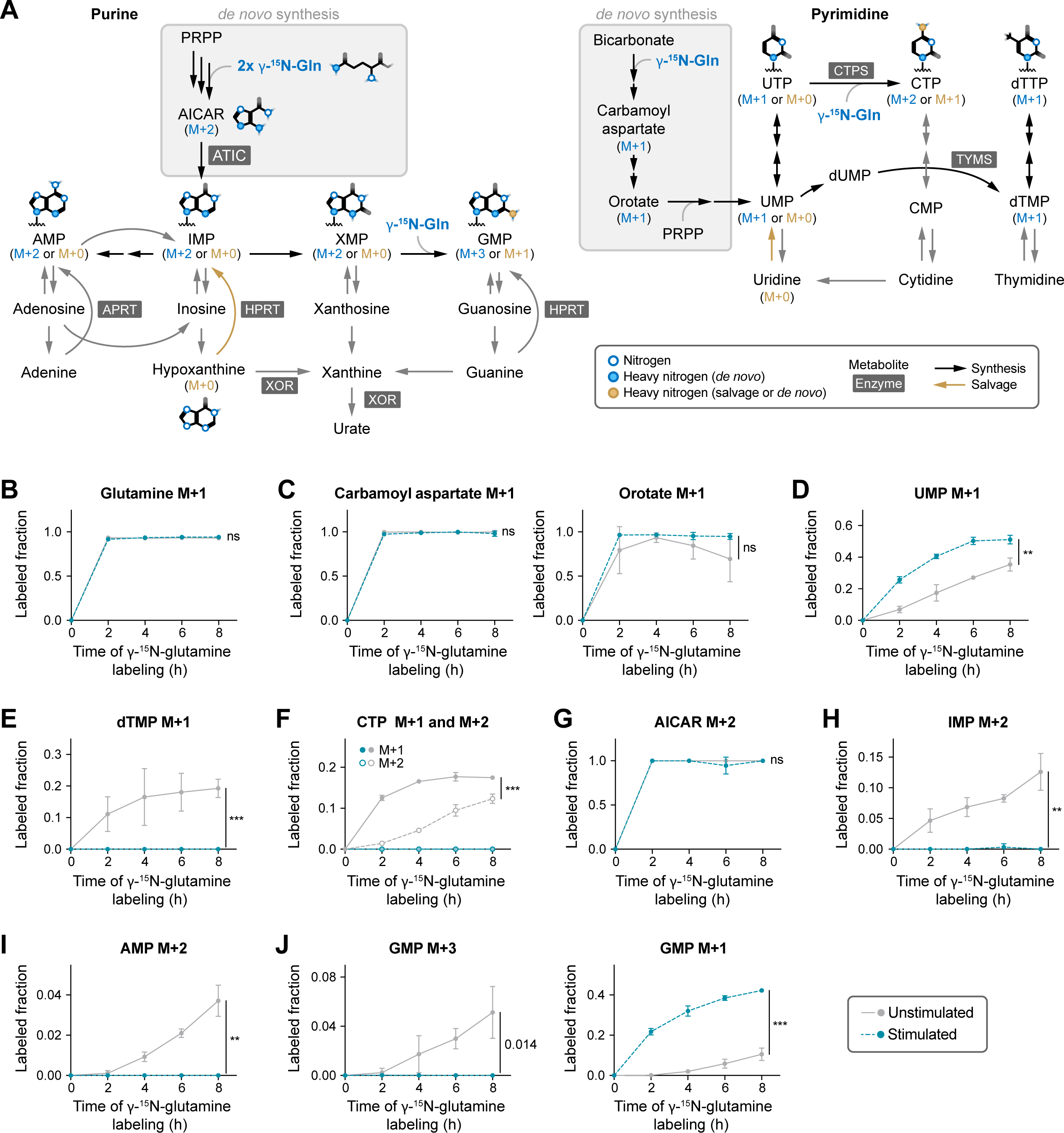
Kinetic glutamine labeling reveals blockages in nucleotide *de novo* synthesis. A) Schematic showing labeling incorporation from γ-^15^N-glutamine into nucleotide metabolism. B–J) Labeling kinetics of indicated intracellular metabolites of nucleotide metabolism in unstimulated or 48h continually stimulated BMDMs. Cells were labeled with γ-^15^N-glutamine for 0–8 hours as indicated on x-axis. Mean ± SD (n=3 independent samples). Statistical analysis was performed with unpaired student’s t test comparing unstimulated and stimulated cells at the 8h time point. ** indicates p<0.01; *** indicates p<0.001; ns indicates not significant (p>0.01).

Labeling rates into pyrimidine nucleotides showed a noticeable difference depending on the stimulation state: labeling into UMP (Figure 2D), and similarly UDP and UTP (Supplemental figure 2A), are slightly faster in stimulated macrophages, while the total levels of UMP are similar (Supplemental figure 2B), suggesting stimulation induces a slightly increased rate in pyrimidine turnover and synthesis up to UMP. In striking contrast, labeling incorporation into nucleotides downstream of UMP, including dTMP (M+1, synthesized via thymidylate synthase (TYMS)), CTP and dCTP (M+1 or M+2, both synthesized via CTP synthase (CTPS)), only occurs significantly in unstimulated macrophages but is completely abolished in stimulated macrophages (Figures 2E–F, Supplemental figure 2C). We further conducted a similar experiment in both unstimulated and 24h stimulated RAW 264.7 macrophages and observed consistent results: intracellular glutamine rapidly became labeled and labeling from γ-^15^N-glutamine was actively incorporated into carbamoyl aspartate, orotate, and UMP in both stimulated and unstimulated macrophages (Supplemental figure 2D, incorporation in unstimulated cells is faster, likely because they are proliferating). However, the further incorporation into dTMP and CTP was only highly active in unstimulated cells, whereas it was completely lost in stimulated cells (Supplemental figure 2E). These results clearly demonstrated that while macrophages in both stimulated and unstimulated states actively perform pyrimidine nucleotide *de novo* synthesis up to uridine nucleotides, the synthesis of cytidine and thymidine nucleotides, through CTPS and TYMS respectively, is shut off upon LPS+IFNγ stimulation.

Similarly, we found drastic changes in purine *de novo* synthesis upon stimulation. 5-Aminoimidazole-4-carboxamide ribonucleotide (AICAR), the last purine *de novo* synthesis intermediate before the synthesis of IMP, rapidly became fully labeled (M+2) from γ-^15^N-glutamine in both stimulated and unstimulated states (Figure 2G). Unstimulated BMDMs continue to incorporate the labeling through purine *de novo* synthesis into IMP (Figure 2H), as well as nucleotides downstream of IMP, e.g., adenine nucleotides (Figure 2I, Supplemental figure 2F). Such labeling is completely lost in stimulated BMDMs. Consistently, in RAW 264.7 macrophages, even though labeling up to AICAR was preserved, labeling into IMP, AMP, and XMP was profoundly inhibited upon stimulation (Supplemental figure 2G). These results demonstrate that the last step of purine *de novo* synthesis, which converts AICAR to IMP, catalyzed by the bi-functional enzyme AICAR transformylase/IMP cyclohydrolase (ATIC), is strongly inhibited upon stimulation.

#### Macrophages switch to nucleotide salvage upon stimulation

While stimulation causes flux through both pyrimidine and purine *de novo* synthesis to be profoundly inhibited, we observed the abundance of most nucleotides, except for dTTP, was largely maintained upon stimulation (Figure 1C), suggesting cells switch to other pathways to obtain nucleotides upon stimulation. Indeed, kinetic tracing with γ-^15^N-glutamine revealed that even though the generation of M+3 GMP through *de novo* synthesis was completely suppressed in stimulated BMDMs, the rate of M+1 GMP production increased substantially (Figure 2J). M+1 GMP is generated when cells salvage unlabeled hypoxanthine, xanthine, or inosine to produce GMP via GMP synthase (GMPS) because only one labeled nitrogen is added during this process (Figure 2A). This suggests stimulated macrophages switch to nucleotide salvage.

We next sought to quantify how the contributions of salvage and *de novo* synthesis to the production of nucleotides change over the timecourse of continual or acute stimulation. To this end, we performed pseudo-steady state γ-^15^N-glutamine tracing in BMDMs stimulated for various durations (cells were always incubated with glutamine tracer for 24h prior to measurement in this experiment). Upon continual stimulation, the fraction of M+1 labeled GMP (again, indicating GMP produced by salvage) increased gradually over time of stimulation; the fraction of M+3 GMP (indicating *de novo* synthesis) decreased upon stimulation to near zero by 48h (Figure 3A). Similarly upon acute stimulation, the fraction of M+1 labeled GMP increased, and the fraction of M+3 GMP decreased to near zero by 48-72h. However, the fraction of M+3 GMP starts to recover at 96h (Figure 3A). Together, these data demonstrate that upon stimulation, cells gradually shift from purine *de novo* synthesis towards salvage. The substantial shift can even be induced by acute stimulation and have a persistent effect for several days, but eventually can recover.

**Figure 3:**
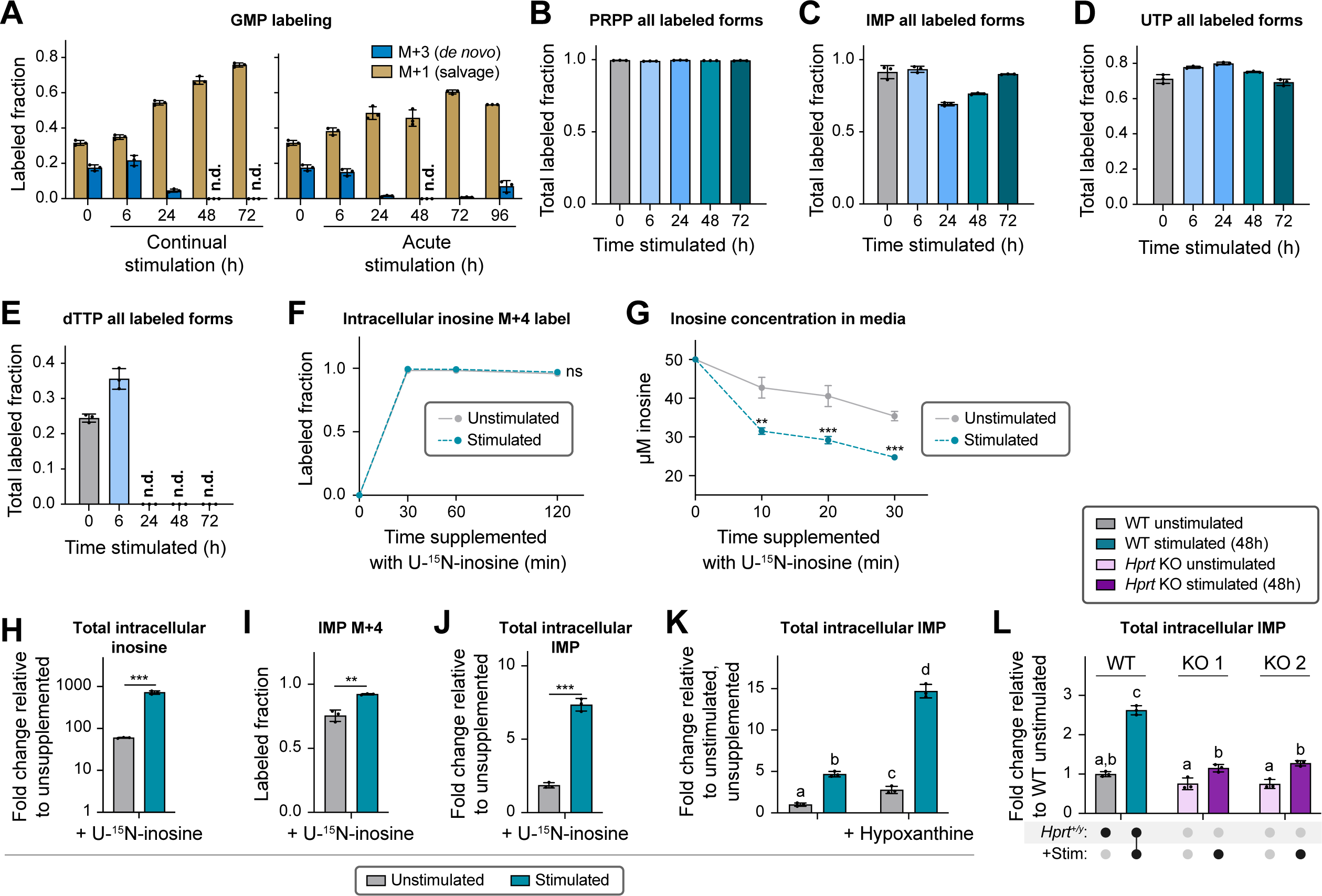
Macrophages shift to salvage for maintaining purines upon stimulation. A) Labeling pattern of M+1 and M+3 GMP in BMDMs over a timecourse of continual (left) or acute (right) stimulation. Cells were labeled with γ-^15^N-glutamine for 24h before analysis. B–E) Total labeled fraction of (B) PRPP, (C) IMP, (D) UTP and (E) dTTP in BMDMs over a timecourse of continual stimulation. Cells were labeled with U-^13^C-glucose for 24h before analysis. F) Labeling kinetics of intracellular inosine (M+4) in unstimulated or stimulated BMDMs. Cells were supplemented with 50 µM U-^15^N-inosine for 0–120 minutes as indicated on x-axis. G) Concentration of inosine remaining in media after unstimulated or stimulated BMDMs were incubated with media supplemented with 50 µM inosine for 0–30 minutes. H) Fold change of intracellular inosine in unstimulated or stimulated BMDMs after supplementing with 50 µM U-^15^N-inosine for 30 minutes, compared to cells of the same stimulation state but without inosine supplementation. I) Labeled fraction of IMP (M+4) in unstimulated or stimulated BMDMs, supplemented with 50 µM U-^15^N-inosine for 30 minutes. J) Fold change of intracellular IMP in unstimulated or stimulated BMDMs after supplementing with 50 µM U-^15^N-inosine for 30 minutes, compared to cells of the same stimulation state but without inosine supplementation. K) Relative abundance of intracellular IMP in unstimulated or stimulated BMDMs, with or without supplementation of 50 µM hypoxanthine for 48 hours. Data are presented relative to abundance in unstimulated, unsupplemented cells. L) Fold change of intracellular IMP in wildtype or *Hprt* KO (two separate clones) RAW 264.7 cells, either unstimulated or stimulated. Data are presented relative to unstimulated wildtype. F–L) Stimulated cells are continually stimulated for 48h. A–L) Mean ± SD (n=3 independent samples). N.d. indicates not detected. F–J) Statistical analysis was performed with unpaired student’s t test comparing unstimulated and stimulated cells. The analysis for F) was performed at the final timepoint. ** indicates p<0.01; *** indicates p<0.001; ns indicates not significant (p>0.01). K–L) Statistical analysis was performed using one-way ANOVA followed by post hoc Tukey’s test. Bars with different lower-case letters (a, b, c, or d) indicate a statistically significant difference with p<0.05.

Both nucleotide *de novo* synthesis and salvage allow macrophages to turn over their nucleotide pools. To examine general nucleotide turnover rate, we also performed tracing with U-^13^C-glucose. U-^13^C-glucose can be incorporated into the ribose moiety of nucleotides through both *de novo* synthesis and the salvage of nitrogenous bases via the common precursor for purine and pyrimidine nucleotides, PRPP. PRPP labeled completely (M+5) from U-^13^C-glucose in unstimulated and stimulated BMDMs (Figure 3B). In general, the labeling of IMP is maintained at a high fraction throughout the stimulation timecourse (Figure 3C). This suggests overall purine nucleotide turnover remained highly active upon stimulation despite *de novo* synthesis of IMP shutting off, pointing to active turnover by salvage instead. Consistent with blocked *de novo* synthesis upon stimulation, we observed the majority of labeled ATP and GTP is M+5 labeled (corresponding to labeling only on the ribose moiety), but higher labeled forms are diminished (Supplemental figure 3A). (These are nucleotides with labeling on the purine ring resulting from *de novo* purine synthesis using precursors synthesized from U-^13^C-glucose, such as serine or glycine.) For pyrimidine nucleotides, the turnover of UTP is maintained (Figure 3D). In sharp contrast, labeling into dTTP is no longer detected after 24h of stimulation (Figure 3E), suggesting stimulated macrophages do not significantly salvage thymine by attaching a new ribose moiety. Furthermore, we found expression of both thymidine kinase 1 and 2, which are required for thymidine salvage, are greatly reduced upon stimulation (Supplemental figure 3B). The lack of both *de novo* synthesis and salvage can cause the unique depletion of dTTP among all nucleotides (Figure 1C).

As the results above showed that stimulated macrophages switch to relying on the salvage pathway to maintain the turnover of purine nucleotides, we next directly assessed the changes in cells’ capability to salvage purine nucleosides and bases upon stimulation. When supplemented with U-^15^N-inosine, BMDMs rapidly take it up from media, and intracellular inosine quickly becomes fully labeled (Figures 3F–G). Compared to unstimulated BMDMs, stimulated BMDMs uptake inosine at a significantly faster rate (Figure 3G) and have a ∼10-fold higher buildup of intracellular inosine (Figure 3H), suggesting stimulated macrophages have a greater potential to utilize extracellular inosine. Both unstimulated and stimulated macrophages can salvage extracellular inosine to produce IMP, as indicated by the generation of labeled IMP from U-^15^N-inosine. Noticeably when supplemented with inosine, both the fraction of labeled IMP and the increase in IMP total abundance are significantly higher in stimulated macrophages (Figures 3I–J), which is consistent with their greater capability to salvage purine nucleosides. Similarly, when cells are supplemented with hypoxanthine, IMP level accumulates, and the accumulation is higher in stimulated macrophages (Figure 3K), showing stimulated macrophages can also very actively salvage purine bases via hypoxanthine-guanine phosphoribosyltransferase (HPRT).

We hypothesized that the increased ability to salvage purine bases (via HPRT or adenine phosphoribosyltransferase (APRT)) and nucleosides (either directly by nucleoside kinases, or indirectly by first breaking the nucleosides down to bases then salvage via HPRT or APRT) is the reason that total intracellular IMP level is increased in stimulated macrophages (Figure 1C, Supplemental figure 1D), despite the fact that IMP *de novo* synthesis activity is shut off. To test this hypothesis, we generated two *Hprt* knockout lines in RAW 264.7 cells, both of which were confirmed by sequencing and remain highly viable (Supplemental figure 3C). As expected, purine salvage upon stimulation is reduced in *Hprt* KO cells, as evidenced by a significantly smaller increase in the M+1 labeled GTP fraction from γ-^15^N-glutamine tracing (Supplemental figure 3D). However, it is not completely abolished, likely because there is still significant expression of *Aprt* (Supplemental figure 3E) and other salvage enzymes. Nonetheless, knocking out *Hprt* nearly completely prevented the accumulation of IMP in stimulated macrophages (Figure 3L), highlighting the importance of salvage via HPRT upon stimulation.

#### Nucleotide degradation is increased but complete oxidation of purine bases is blocked upon stimulation

The metabolomic analysis highlighted that some of the most significant metabolomic changes upon stimulation are the buildup of nucleosides and nitrogenous bases, which result from nucleotide degradation and can serve as substrates for salvage (Figure 1C). Moreover, the RNAseq analysis revealed several important nucleotide degradation enzymes, including *Pnp1*, *Pnp2*, and *Upp1*, increased substantially upon stimulation (Figure 1E). Therefore, we hypothesized that nucleotide degradation is significantly rewired in response to LPS+IFNγ stimulation. To probe nucleotide degradation, we first performed kinetic U-^13^C-glucose tracing. U-^13^C-glucose is incorporated into nucleotides like IMP quickly in both unstimulated and stimulated BMDMs (Supplemental figure 4A). The kinetic delay of inosine labeling compared to IMP labeling indicates IMP degradation activity. We found that compared to unstimulated BMDMs, stimulated BMDMs incorporate labeling from IMP to inosine much faster (Supplemental figure 4B), suggesting increased IMP degradation, which can explain the observed accumulation of inosine upon stimulation.

Nucleosides, like inosine, can be further broken down to corresponding bases, and both nucleosides and bases can be released extracellularly by macrophages (Halbrook et al., 2019). We found that not only did the intracellular levels of nucleosides and nitrogenous bases accumulate upon stimulation (Figure 1C), the release of most detected nucleosides and bases into the media also increased greatly with stimulation (Supplemental figure 4C). These results, consistent with increased *Pnp* and *Upp* expression, suggest that in addition to increased nucleotide degradation to nucleosides, the degradation of nucleosides to nitrogenous bases likely also increased upon stimulation.

To directly measure purine nucleoside degradation capability, we supplemented stimulated (48h) or unstimulated BMDMs with U-^15^N-inosine and followed the metabolic fate of inosine. In both stimulated and unstimulated states, macrophages rapidly convert the inosine to hypoxanthine, which became nearly fully labeled (M+4) and accumulated greatly within 30 minutes (Figure 4A). Noticeably, stimulated macrophages build up ∼5-fold more hypoxanthine from inosine supplementation, consistent with stimulation increasing inosine degradation activity via PNP. The hypoxanthine generated was further released into media, with a release rate 2-3-fold faster in stimulated BMDMs (Figure 4B). Hypoxanthine can be further degraded to xanthine and then urate via the action of xanthine oxidoreductase (XOR). In striking contrast to the finding that stimulated macrophages generate more hypoxanthine from U-^15^N-inosine, we found stimulated macrophages accumulated much less labeled xanthine (M+4) from inosine supplementation— this despite a higher baseline xanthine level (M+0) (Figure 4C). The release of inosine-derived xanthine into media is much slower in stimulated macrophages as well (Figure 4D). Even more striking differences were observed in the final degradation product urate— inosine supplementation resulted in urate released into media by unstimulated but not stimulated macrophages (Figure 4E) and a much lower total intracellular urate level in stimulated cells compared to unstimulated cells (Figure 4F). Together, these results revealed that an increase in PNP activity upon stimulation supports increased hypoxanthine production, but the activity of XOR to completely degrade hypoxanthine to urate is strongly inhibited. This inhibition can contribute to the observed accumulation of hypoxanthine, diverting flux away from its complete degradation, and instead into its salvage to IMP via HPRT to maintain purine nucleotides.

**Figure 4:**
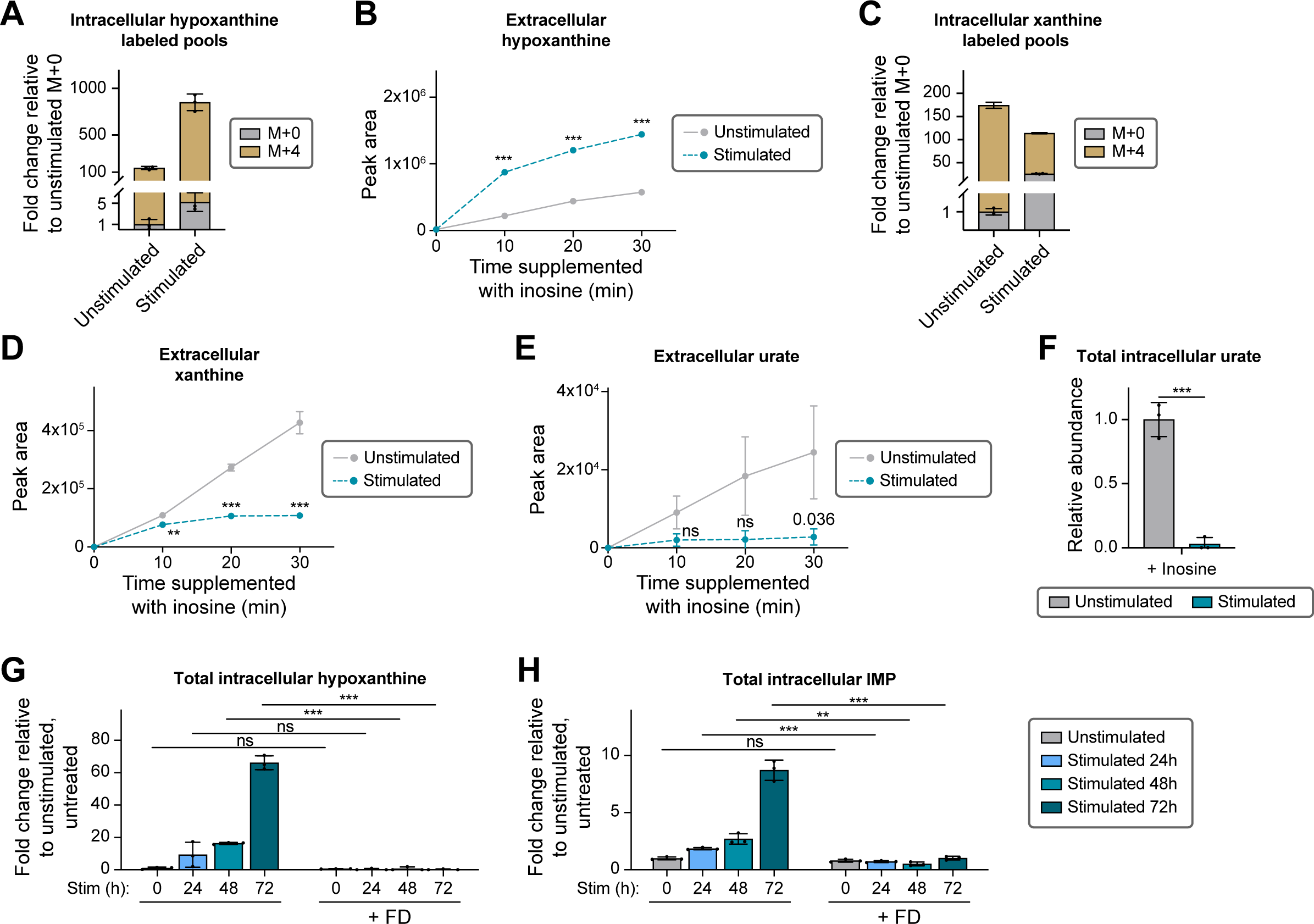
Alterations in nucleotide degradation upon stimulation. A) Relative intracellular abundance of M+0 and M+4 hypoxanthine in unstimulated or stimulated BMDMs supplemented with 50 µM U-^15^N-inosine for 30 minutes. Data are presented relative to the level of M+0 hypoxanthine in unstimulated BMDMs. B) Release of hypoxanthine into media over time by unstimulated or stimulated BMDMs, supplemented with 50 µM inosine. C) Relative intracellular abundance of M+0 and M+4 xanthine in unstimulated or stimulated BMDMs supplemented with 50 µM U-^15^N-inosine for 30 minutes. Data are presented relative to the level of M+0 xanthine in unstimulated BMDMs. D–E) Release of (D) xanthine and (E) urate into media over time by unstimulated or stimulated BMDMs, supplemented with 50 µM inosine. F) Relative intracellular abundance of urate in unstimulated or stimulated BMDMs, supplemented with 50 µM inosine for 10 minutes. Data are presented relative to abundance in unstimulated cells. A–F) Stimulated BMDMs are continually stimulated for 48h. G–H) Relative intracellular abundance of (G) hypoxanthine, and (H) IMP in BMDMs over a timecourse of continual stimulation, with or without treatment of 10 µM forodesine (FD) throughout the timecourse. Data are presented relative to abundance in unstimulated, untreated cells. A–H) Mean ± SD (n=3 independent samples). B, D–F) Statistical analysis was performed with unpaired student’s t test comparing unstimulated to stimulated cells at each time point. G–H) Statistical analysis was performed with unpaired student’s t test comparing untreated to FD treated cells at each time point. ** indicates p<0.01; *** indicates p<0.001; ns indicates not significant (p>0.01).

To further validate the role of PNP in generating hypoxanthine and providing substrate for salvage, we treated macrophages with the PNP inhibitor forodesine. As expected, forodesine treatment caused the accumulation of PNP’s substrate, inosine (Supplemental figure 4D), and completely prevented the stimulation-induced accumulation of hypoxanthine (Figure 4G), confirming hypoxanthine accumulation resulted from increased PNP-dependent nucleoside degradation. Furthermore, forodesine treatment completely prevented the stimulation-induced accumulation of IMP (Figure 4H). This agrees with the findings in *Hprt* KO cells (Figure 3L), demonstrating salvage of PNP-produced hypoxanthine via HPRT is the main source of IMP buildup in stimulated macrophages. Consistently, by limiting hypoxanthine availability, PNP inhibition significantly reduced the fraction of GMP that resulted from salvage, as indicated by a lower M+1 fraction from γ-^15^N-glutamine tracing (Supplemental figure 4E), and caused further buildup of the other HPRT substrate, PRPP, in stimulated macrophages (Supplemental figure 4F).

### Mechanisms driving the rewiring of nucleotide metabolism

The studies above revealed systematic rewiring of nucleotide metabolism in macrophages upon LPS+IFNγ stimulation and identified a series of important regulation points, as summarized in Figure 5A. These include: (1) *De novo* synthesis of purine nucleotides is profoundly inhibited, and the blockage particularly occurs at the step of IMP synthesis from AICAR, catalyzed by ATIC; (2) *De novo* synthesis of pyrimidine nucleotides is also profoundly inhibited, and the blockage occurs at CTP and dTMP production from UMP, catalyzed by CTPS and TYMS, respectively; (3) Cells switch to relying on salvage to maintain most nucleotides; (4) Nucleotide degradation to nucleosides and nitrogenous bases, catalyzed by PNP and UPP, is generally increased, but full degradation of purine bases through the reaction catalyzed by XOR is inhibited, shunting flux into purine salvage. We next sought to elucidate the molecular mechanisms driving these key changes in nucleotide metabolism reactions.

**Figure 5:**
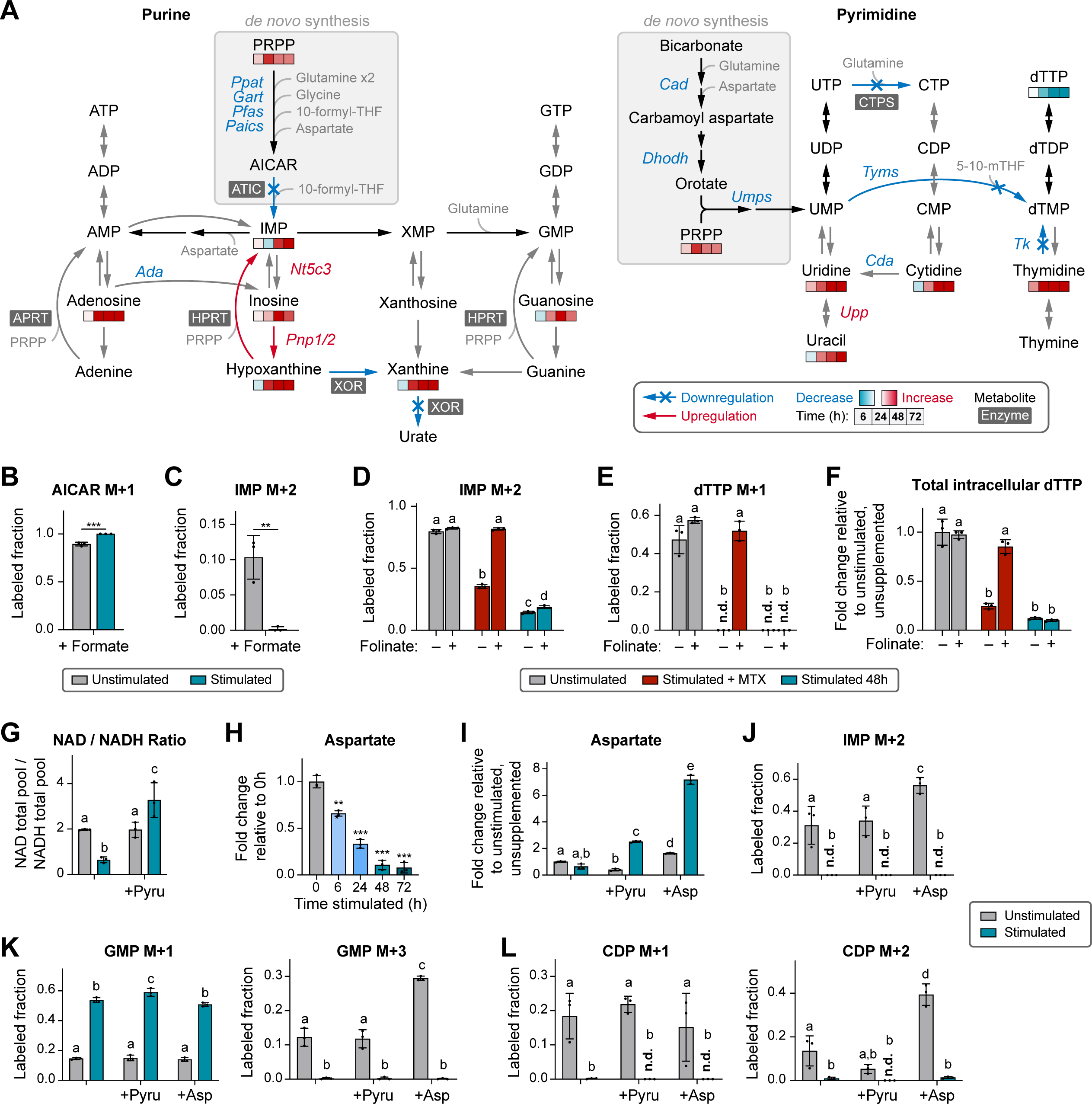
The inhibition of nucleotide *de novo* synthesis is not driven by changes in the folate pathway, redox state, or aspartate level. A) Summary of nucleotide metabolism rewiring in macrophages upon stimulation. Changes in metabolite levels over time are presented by color code in heatmap bars compared to unstimulated cells. All the significant changes in transcript level 24h after acute stimulation revealed by transcriptomics are presented as color codes of the gene name. Blockages of reaction activity are indicated by blue arrows with an X and increases of reaction activity are indicated by red arrows. B–C) Labeled fraction of (B) AICAR and (C) IMP in unstimulated or stimulated BMDMs supplemented with 1mM ^13^C-formate for 24h. D–F) Labeled fraction of (D) IMP and (E) dTTP, and (F) relative intracellular dTTP abundance in RAW 264.7 cells that are unstimulated, stimulated, or unstimulated and treated with 200nM methotrexate (MTX), labeled with γ-^15^N-glutamine for 24h. Cells were additionally supplemented with or without 1 µM folinate throughout the timecourse. G) Intracellular NAD to NADH ratio in unstimulated or stimulated BMDMs with or without supplementation of 2.5mM pyruvate. H) Relative intracellular abundance of aspartate in BMDMs over a timecourse of continual stimulation. I) Relative intracellular abundance of aspartate in unstimulated or stimulated BMDMs, with or without supplementation of 2.5mM pyruvate or 5mM aspartate throughout the timecourse. Data are presented relative to abundance in unstimulated, unsupplemented cells. J–L) Fraction of (J) M+2 labeled IMP, (K) M+1 and M+3 labeled GMP, and (L) M+1 and M+2 labeled CDP, after 24h γ-^15^N-glutamine labeling in unstimulated or stimulated BMDMs, with or without supplementation of 2.5mM pyruvate or 5mM aspartate throughout the timecourse. N.d. indicates not detected. B–G, I–L) Stimulated cells are continually stimulated for 48h. B–L) Mean ± SD (n=3 independent samples). B–C) Statistical analysis was performed with unpaired student’s t test comparing unstimulated to stimulated cells. H) Statistical analysis was performed using one-way ANOVA followed by post hoc Dunnett’s test comparing all groups to unstimulated cells (0h). ** indicates p<0.01; *** indicates p<0.001. D–G, I–L) Statistical analysis was performed using one-way ANOVA followed by post hoc Tukey’s test. Bars with different lower-case letters (a, b, c, d, or e) indicate a statistically significant difference with p<0.05.

#### One-carbon metabolism supplements do not rescue the inhibition of nucleotide de novo synthesis

Both the reactions catalyzed by ATIC and TYMS require one-carbon units supplied by the folate pathway (10-formyl-THF and 5,10-methylene-THF, respectively) as substrates to support the production of IMP and dTMP. Moreover, transcriptomic analysis revealed stimulation causes a significant reduction in the expression of some folate pathway enzymes (*Shmt1*, *Shmt2*, *Dhfr*, *Mthfd1*, etc.) and the rate-limiting enzyme in the serine *de novo* synthesis pathway, *Phgdh*, which feeds into one-carbon metabolism (Supplemental figure 5A). Therefore, we tested the possibility that a lack of one-carbon units causes the reduced flux through ATIC and TYMS.

First, we supplemented BMDMs with 1mM ^13^C-formate, an established method to increase the supply of one-carbon units (Momb et al., 2013; Labuschagne et al., 2014; Ducker et al., 2016). With this supplement, AICAR became nearly fully labeled (M+1) in both unstimulated and 48h stimulated cells (Figure 5B), demonstrating ^13^C-formate is sufficiently converted into available one-carbon units supplying AICAR synthesis. In unstimulated macrophages, the action of ATIC further adds another labeled carbon from 10-formyl-THF to AICAR, generating M+2 labeled IMP (Figure 5C), as well as labeled downstream nucleotides such as ATP and GTP (Supplemental figures 5B–C). However, M+2 labeled IMP, ATP, or GTP was completely absent in stimulated cells (Figure 5C, Supplemental figures 5B–C), indicating that formate supplementation did not rescue the blockage at ATIC upon stimulation. Nor did it rescue dTTP synthesis— no labeled dTTP was detected from ^13^C-formate, and the dTTP depletion upon stimulation persisted despite formate supplementation (Supplemental figure 5D).

It is possible that insufficient levels of the THF cofactor, rather than one-carbon units (which would be supplied by formate), can limit one-carbon metabolism and thus further limit ATIC and TYMS activities. To test this possibility, we supplemented RAW 264.7 cells with folinate, which has been shown effective in rescuing limited folate availability, such as that caused by the treatment with the anti-folate drug methotrexate (Iqbal et al., 2001; Howard et al., 2016). Indeed, treating unstimulated cells with methotrexate to inhibit folate metabolism can strongly inhibit IMP and dTTP *de novo* synthesis, as evidenced here by the substantially decreased labeling into these compounds using the γ-^15^N-glutamine tracing assay (Figures 5D–E), providing a positive control. However, there is an important difference between methotrexate-induced and stimulation-induced inhibition of nucleotide *de novo* synthesis: while the former can be fully rescued by folinate supplementation, stimulation-induced inhibition is not affected by folinate (Figures 5D–E). Consequentially, the depletion of dTTP caused by methotrexate treatment can be rescued by folinate supplementation, but the depletion of dTTP induced by stimulation cannot (Figure 5F). Together, these experiments show strong evidence that mechanisms other than insufficiency of THF or one-carbon units are primarily responsible for the inhibition of nucleotide *de novo* synthesis upon macrophage stimulation.

#### Changes in nucleotide synthesis upon stimulation is largely independent of changing NAD/NADH ratio and aspartate level

When macrophages are stimulated with LPS+IFNγ, respiration rate (Supplemental figure 5E) and citrate cycle flux reduce (Tannahill et al., 2013; Jha et al., 2015; Seim et al., 2019; Ryan and O’Neill, 2020). Related to these changes, we observed the NAD/NADH ratio decreased significantly (Figure 5G), and aspartate, which is synthesized from a citrate cycle intermediate, decreased gradually over the continual stimulation timecourse (Figure 5H). Several previous studies have demonstrated that a decreased NAD/NADH redox ratio can limit the synthesis of nucleotides in cancer cells, because nucleotide synthesis requires electron acceptors to support the synthesis of various precursors, including aspartate (Sulivan et al., 2015; Sulivan et al., 2018; Diehl et al., 2019). To test whether redox stress or the decrease in aspartate are limiting factors for nucleotide synthesis in stimulated macrophages, we supplemented cells with pyruvate, which can act as an electron acceptor, or aspartate— perturbations that have been shown to effectively rescue nucleotide synthesis constrained by a decreased NAD/NADH ratio in cancer cells. We found that pyruvate supplementation effectively rescued the decrease in cellular NAD/NADH ratio in stimulated macrophages (Figure 5G), and both pyruvate and aspartate supplementation completely reversed the decrease of aspartate level in stimulated macrophages (Figure 5I), indicating the success of these supplementations. However, neither pyruvate nor aspartate rescued the inhibition of purine *de novo* synthesis in stimulated macrophages, as indicated by the persistent complete absence of 2-labeled IMP or AMP when tracing with γ-^15^N-glutamine (Figure 5J, Supplemental figure 5F). Purine nucleotides continue to be mainly supplied by salvage upon stimulation, as indicated by an elevated M+1 fraction of GMP and a lack of M+3 fraction (Figure 5K). Similarly, neither pyruvate nor aspartate supplementation rescued the inhibition of CDP synthesis in stimulated macrophages, as indicated by the complete absence of both M+1 and M+2 fractions of CDP from γ-^15^N-glutamine labeling (Figure 5L), despite UMP being labeled (Supplemental figure 5G). These results are consistent with our finding that the inhibition of pyrimidine synthesis does not occur upstream of UMP synthesis, where aspartate is required as a substrate, but rather happens at the downstream steps of CTPS and TYMS (Figures 2D–F). Together, these results showed that the inhibition of *de novo* purine and pyrimidine synthesis upon stimulation is largely independent of changes in NAD/NADH ratio or aspartate level.

#### Nitric oxide-driven inhibition of ATIC and CTPS together with transcriptional changes drive inhibition of nucleotide synthesis

To identify the regulators that may be responsible for the rewiring of nucleotide metabolism, we searched for changes that temporally correlated with the changes in nucleotide metabolism activities. We found that: (1) The inhibition of nucleotide *de novo* synthesis activity occurs soon after the strong induction of inducible nitric oxide synthase (iNOS, encoded by the gene *Nos2*), which macrophages express upon classical activation to produce nitric oxide (NO) for pathogen killing (Figure 6A, Supplemental figure 1F). (2) Upon stimulation, the expression of many nucleotide *de novo* synthesis genes decreases significantly prior to the decrease of nucleotide *de novo* synthesis activity (Figure 1E). Most noticeably, *Tyms* expression decreases substantially upon both acute stimulation (Figure 1E) and continual stimulation (Figure 6B). Therefore, we investigated the role of NO and transcriptional changes in driving the rewiring of nucleotide metabolism.

**Figure 6:**
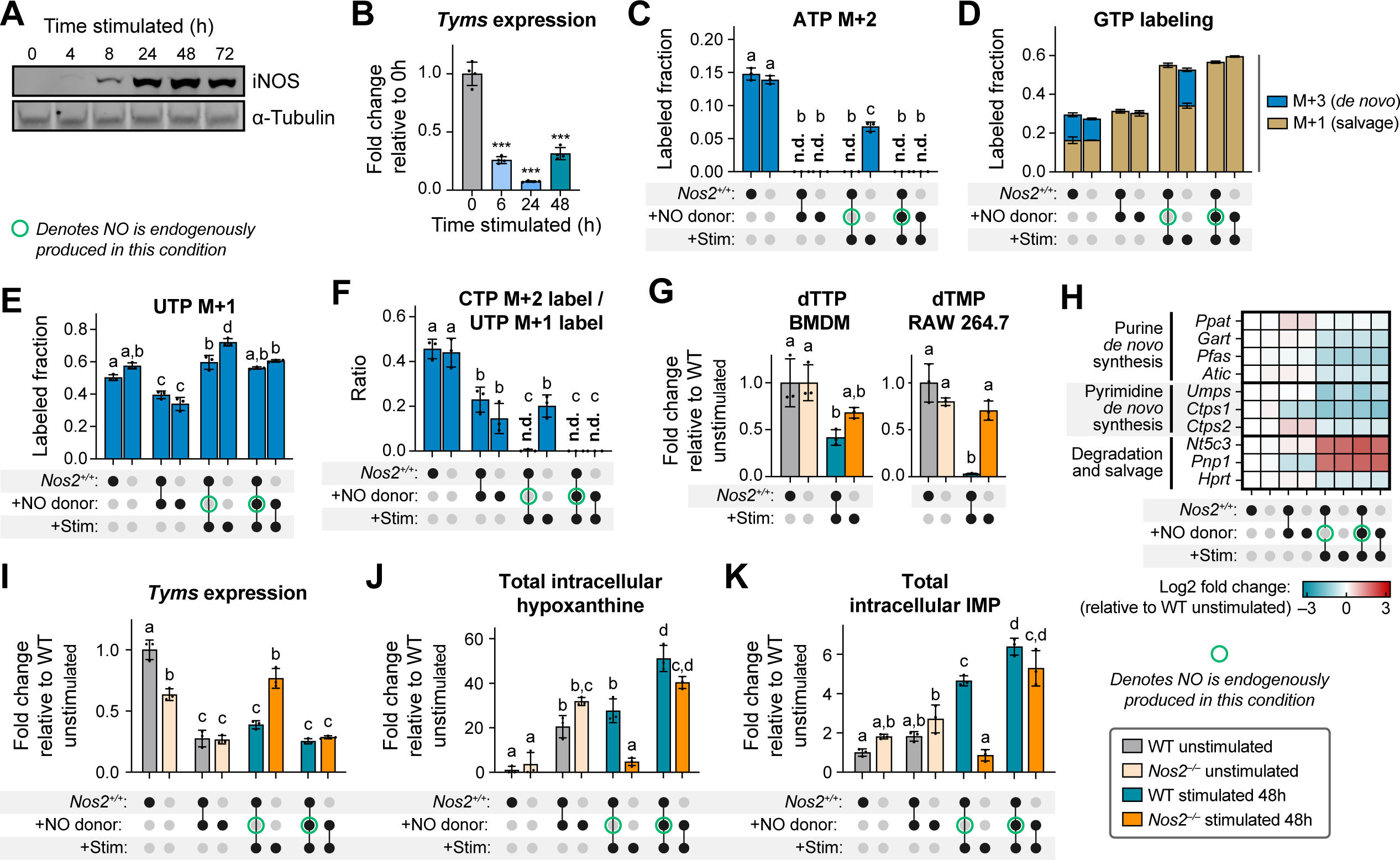
NO-dependent inhibition of ATIC, CTPS and XOR together with transcriptional changes drive the rewiring of nucleotide metabolism. A) iNOS protein levels in BMDMs over a timecourse of continual stimulation. B) Fold change of *Tyms* expression (relative to unstimulated) in BMDMs over a timecourse of continual stimulation. C–F) Labeled fraction of (C) ATP (M+2), (D) GTP (M+1 and M+3), (E) UTP (M+1), and (F) the ratio of labeled CTP (M+2) to labeled UTP (M+1), after 24h γ-^15^N-glutamine labeling in unstimulated or stimulated wildtype or *Nos2*^-/-^ BMDMs, with or without treatment of 200 µM NO donor DETA-NONOate throughout the timecourse. N.d. indicates not detected. G) Left: Relative intracellular dTTP abundance in unstimulated or stimulated wildtype or *Nos2*^-/-^ BMDMs. Right: Relative intracellular dTMP abundance in unstimulated or stimulated wildtype or *Nos2*^-/-^ RAW 264.7 cells. Data are expressed relative to abundance in wildtype unstimulated cells. H) Expression levels of selected genes in purine and pyrimidine *de novo* synthesis, degradation, and salvage pathways, probed by qPCR in unstimulated or stimulated wildtype or *Nos2*^-/-^ BMDMs, with or without treatment of 200 µM NO donor DETA-NONOate throughout the timecourse. Data are presented relative to expression in unstimulated, untreated wildtype cells and displayed on a log2 scale as a heatmap. Each box represents the mean of n=3 independent samples. I–K) Relative *Tyms* expression and relative intracellular abundance of (J) hypoxanthine, and (K) IMP in unstimulated or stimulated wildtype or *Nos2*^-/-^ BMDMs, with or without treatment of 200 µM NO donor DETA-NONOate throughout the timecourse. Data are presented relative to expression in unstimulated, untreated wildtype cells. B–G, I–K) Mean ± SD (n=3 independent samples). C–K) Stimulated cells are continually stimulated for 48h. B) Statistical analysis was performed using one-way ANOVA followed by post hoc Dunnett’s test comparing all groups to unstimulated cells (0h). *** indicates p<0.001. C, E–G, I–K) Statistical analysis was performed using one-way ANOVA followed by post hoc Tukey’s test. Bars with different lower-case letters (a, b, c, or d) indicate a statistically significant difference with p<0.05.

To perturb NO level in macrophages, we took two approaches. First, to limit NO accumulation in macrophages upon stimulation, we used BMDMs isolated from *Nos2*^-/-^ mice, which do not produce NO upon stimulation (Seim et al., 2023). The efficacy is confirmed by the lack of citrulline (the other product of iNOS) accumulation over time after stimulation (Supplemental figure 6A). Second, to increase cellular NO level, we treated unstimulated macrophages or *Nos2*^-/-^ macrophages with the NO donor DETA NONOate to recapitulate what occurs upon stimulation in wildtype macrophages. To examine nucleotide *de novo* synthesis and salvage activity, we performed pseudo-steady state γ-^15^N-glutamine tracing, as illustrated in figure 2A. Across all the conditions, intracellular glutamine remained nearly fully labeled (Supplemental figure 6B). Strikingly, the loss of 2-labeled ATP (Figure 6C) and 3-labeled GTP (Figure 6D) in stimulated wildtype BMDMs, which indicates the blockage of purine *de novo* synthesis, was rescued in stimulated *Nos2*^-/-^ BMDMs. Conversely, treating unstimulated macrophages with NO donor is sufficient to cause the complete loss of *de novo* purine synthesis, like that observed in stimulated macrophages (as indicated by the loss of 2-labeled ATP and 3-labeled GTP). Moreover, treating *Nos2*^-/-^ macrophages with NO donor reversed the rescue of purine *de novo* synthesis upon stimulation (Figures 6C–D). These results show strong evidence that the shutdown of purine *de novo* synthesis upon stimulation is driven by NO production.

Mirroring the effects on purine *de novo* synthesis, we found the contribution of salvage to GTP production, as indicated by the fraction of M+1 labeled GTP from γ-^15^N-glutamine, increases greatly in conditions where cellular NO level is high (in wildtype cells upon stimulation or in cells treated with NO donor). This finding demonstrates that NO drives the switch from *de novo* synthesis to salvage for purine turnover. Interestingly, overall GTP labeling turnover (i.e., *de novo*-derived M+3 and salvaged-derived M+1 combined) increases upon stimulation. The stimulation-induced increase is independent of NO status, as it stayed the same regardless of *Nos2* genotype or NO donor treatment (Figure 6D).

As for pyrimidine *de novo* synthesis, pseudo-steady state γ-^15^N-glutamine tracing showed that UTP remained ∼50% M+1 labeled in all conditions, suggesting pyrimidine *de novo* synthesis up to UTP is largely not altered by NO, while the labeling is slightly higher in all stimulated conditions (Figure 6E). We have demonstrated above that upon stimulation, pyrimidine *de novo* synthesis is maintained up to UTP, but blocked at the steps of CTP and dTMP synthesis. The synthesis of CTP from UTP by CTPS is indicated by the conversion of M+1 labeled UTP to M+2 labeled CTP (Figure 2A). We found the loss of M+2 labeled CTP upon stimulation is partially rescued in *Nos2*^-/-^ BMDMs (Figure 6F). Furthermore, treatment with NO donor significantly reduced the ratio of M+2 CTP to M+1 UTP in unstimulated macrophages, partially recapitulating the decrease of CTPS flux upon stimulation. Additionally, treating stimulated *Nos2*^-/-^ macrophages with NO donor reversed the rescue of CTP synthesis (Figure 6F). These results demonstrate that NO is a significant regulator in driving the blockage of CTPS upon stimulation. *Nos2*^-/-^ also partially rescued the depletion of dTTP upon stimulation in BMDMs, and to a greater extent, rescued the dTMP depletion in stimulated RAW 264.7 cells (Figure 6G), suggesting NO plays a significant role in inhibiting dTMP synthesis.

As the results above provided strong evidence that NO is a key driver in the inhibition of nucleotide *de novo* synthesis, we next examined whether NO’s effect occurs through, or independent of, transcriptional changes. Consistent with the transcriptomic analysis upon acute stimulation, qPCR analysis confirmed the expression of many purine *de novo* synthesis genes decreased upon continual stimulation. We found these changes are induced by stimulation in a NO-independent manner, as *Nos2*^-/-^ or treatment with NO donor had no major consistent effect on their expression (Figure 6H). The decrease of these genes, such as *Gart* and *Pfas,* is unlikely to be strong enough to cause the complete block of overall purine *de novo* synthesis flux upon stimulation, because we showed flux is mostly maintained up to AICAR production but is shut off at the ATIC step. Furthermore, *Nos2*^-/-^ can nearly completely rescue the inhibition of purine *de novo* synthesis in stimulated cells despite it not rescuing the expression of these upstream purine *de novo* synthesis genes. This suggests that NO-dependent inhibition of ATIC activity is the main driver of purine *de novo* synthesis inhibition, although the NO-independent transcriptional downregulation upon stimulation may have a minor role in augmenting the inhibition, particularly at later time points. Consistent with this suggestion, in *Nos2*^-/-^ cells, AMP labeling from glutamine is slightly increased 24h after stimulation, which likely resulted from increased overall purine turnover, but then slightly decreased at 48h (Supplemental figure 6C), when the NO-independent transcriptional downregulation in purine *de novo* synthesis enzymes is likely to take effect. Moreover, the combined action of NO-dependent ATIC inhibition and NO-independent transcriptional downregulation can explain the very interesting dynamics we observed in AICAR level: an initial strong accumulation followed by a decrease over a timecourse of continual stimulation (Supplemental figure 6D). The early peak is likely due to the quick inhibition of ATIC after NO production is induced, cutting off AICAR consumption flux, while the later AICAR decrease can be caused by transcriptional downregulation of upstream enzymes taking effect at this time, limiting the production of AICAR (Supplemental figure 6D).

With respect to pyrimidine *de novo* synthesis, we found both *Ctps1* and *Ctps2* expression decreased significantly upon stimulation (Figure 6H), and this decrease is largely NO independent, as the expression does not vary in correlation with NO level across *Nos2* genotypes and NO donor treatment (Figure 6H). Such NO-independent decreases in *Ctps1* and *Ctps2* expression can act together with the NO-dependent inhibition of CTPS activity to block CTP synthesis flux from UTP upon stimulation, explaining the portion of CTP synthesis inhibition that is not fully rescued in stimulated *Nos2*^-/-^ BMDMs (Figure 6F). Interestingly, the transcriptional downregulation of *Tyms* in stimulated macrophages can be significantly rescued by *Nos2*^-/-^(Figure 6I). Conversely, NO donor treatment decreased *Tyms* expression in unstimulated macrophages, fully recapitulating the stimulation-induced effect, and NO donor treatment reversed the rescue of *Tyms* expression in stimulated *Nos2*^-/-^macrophages. These results show that NO production drives the transcriptional *Tyms* downregulation, which in turn drives the inhibition of dTTP synthesis and dTTP depletion upon stimulation.

#### NO-driven inhibition of XOR diverts flux to purine nucleotide salvage

Finally, we found the key regulation points in nucleotide degradation are also regulated by a combination of NO-independent transcriptional changes and NO-dependent inhibition. We showed above that upon stimulation, nucleotide degradation to nitrogenous bases is increased. Indeed, nucleotide degradation genes such as *Nt5c3* and *Pnp* increase expression by several folds upon stimulation, and we found this increase is not affected by *Nos2*^-/-^ (Figure 6H), suggesting NO-independent transcriptional upregulation contributes to increased nucleotide degradation.

We also showed that downstream of PNP, the complete degradation of the purine base hypoxanthine by XOR is inhibited upon stimulation, allowing cells to greatly accumulate intracellular hypoxanthine and divert flux into purine salvage (Figure 5A). We found that NO plays a critical role in such XOR inhibition and re-distributing the metabolic fate of purine bases. Knocking out *Nos2* completely prevented the accumulation of hypoxanthine upon stimulation (Figure 6J), and similarly prevented the accumulation of IMP upon stimulation (Figure 6K), which, as we showed above, mainly results from increased hypoxanthine salvage. These effects can be reversed by treating *Nos2^-/-^* BMDMs with NO donor (Figures 6J–K). Moreover, treating unstimulated macrophages with NO donor is sufficient to cause the accumulation of hypoxanthine and IMP, recapitulating the changes induced by stimulation. These results provide evidence that NO drives the inhibition of XOR and diverts flux into purine salvage upon stimulation. In further confirmation of this finding, we observed that in *Nos2*^-/-^ BMDMs, xanthine never accumulated like in wildtype cells (Supplemental figure 6E). Additionally, PRPP increases over time after stimulation to a greater extent in *Nos2*^-/-^ cells (Supplemental figure 6F), which is consistent with *Nos2*^-/-^ cells having reduced PRPP consumption via purine salvage due to a lack of xanthine and hypoxanthine as salvage substrates.

### Alteration of nucleotide metabolism impacts macrophage function and pathogen growth

The research above revealed the substantial rewiring of nucleotide metabolism upon stimulation and its underlying mechanisms. Given the remarkable switch that macrophages make upon stimulation to relying on salvage instead of *de novo* synthesis for maintaining purine nucleotides, we next investigated the significance of such a switch on macrophage functions.

#### Purine salvage by HPRT is required for efficient macrophage migration

To understand the functional importance of purine salvage, we first performed RNA-seq analysis in unstimulated and stimulated *Hprt* KO macrophages and compared the transcriptomic profiles to their wildtype counterparts. *Hprt* KO had a greater impact on global gene expression (causing more significant changes) in the stimulated state compared to the unstimulated state (Supplemental figures 7A–B), consistent with cells increasing their reliance on HPRT-dependent salvage upon stimulation. We analyzed the GO enrichment of differentially expressed genes in *Hprt* KO compared to wildtype macrophages. The top enriched process that is downregulated in *Hprt* KO cells (in both the unstimulated state, and more profoundly, in the stimulated state) is cell migration (Figure 7A), which is essential for macrophages’ function in innate immunity. Upon sensing signals associated with infection such as LPS, macrophages migrate to the site of infection and remodel the tissue using enzymes such as matrix metalloproteinases. Given that *Hprt* KO reduces the stimulation-induced upregulation of many genes involved in chemotaxis and extracellular matrix remodeling, we sought to specifically validate the impact of purine salvage in macrophage migration and invasion. Wildtype or *Hprt* KO macrophages were seeded in transwells coated with Matrigel and migration was induced by LPS in the bottom chamber (Supplemental figure 7C). Consistent with the transcriptional changes, *Hprt* KO macrophages showed substantially reduced transwell migration towards LPS (Figure 7B), and likewise reduced migration towards another chemoattractant C5a (Supplemental figure 7D). These results demonstrate that purine salvage is important for stimulation-induced macrophage migration.

**Figure 7:**
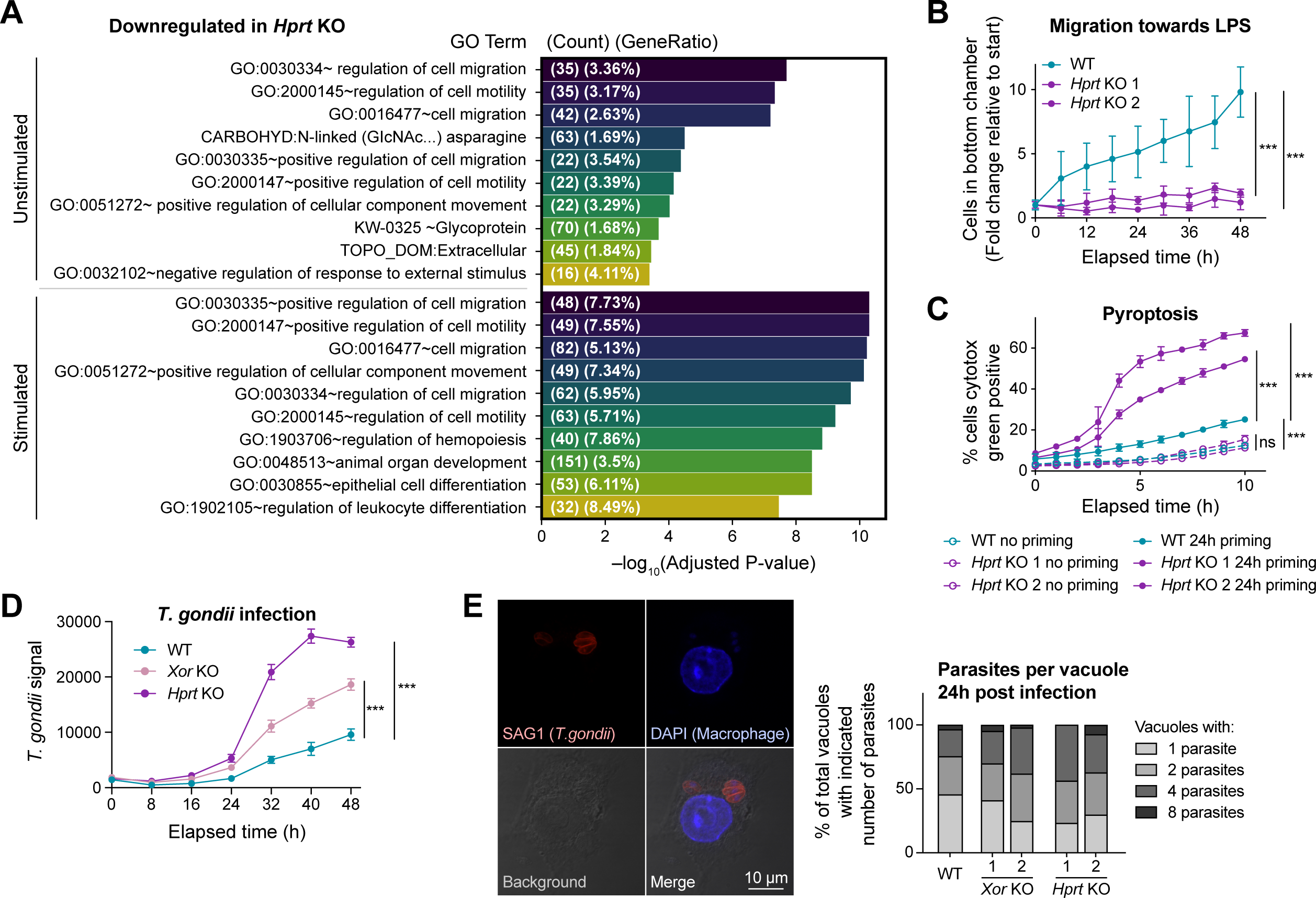
Alteration of nucleotide metabolism impacts macrophage function and pathogen growth. A) Gene ontology enrichment of the pathways that are most downregulated in *Hprt* KO macrophages compared to wildtype. B) Migration of wildtype and *Hprt* KO (2 clones) RAW 264.7 cells across a Matrigel-coated transwell towards LPS. Mean ± SD (n=5 independent samples). C) Percentage of wildtype or *Hprt* KO (2 clones) RAW 264.7 cells that are Cytotox green positive at indicated time points after treatment with 10 µM nigericin. Cells were either primed with LPS+IFNγ for 24h (closed circle) or not primed (open circles). Mean ± SD (n=3 independent samples). D) Total fluorescence signal from *T. gondii* at indicated time points after wildtype, *Xor* KO, or *Hprt* KO RAW 264.7 macrophages were infected with mCherry ME49 *T. gondii*. Mean ± SD (n=4 independent samples). E) Wildtype, *Xor* KO, or *Hprt* KO RAW 264.7 macrophages were infected with mCherry ME49 *T. gondii* for 24h, then fixed and incubated with primary antibody against *T. gondii* surface antigen 1 (SAG1) and counterstained with DAPI. Left: Representative immunofluorescence image of wildtype cell infected by ME49 *T. gondii*. Right: percentage of vacuoles with 1, 2, 4, or 8 parasites per vacuole, quantified from 10 random fields of view per coverslip. B–D) Statistical analysis was performed using one-way ANOVA followed by post hoc Dunnett’s test. *** indicates p<0.001; ns indicates not significant (p>0.01).

#### Loss of HPRT-dependent purine salvage increases pyroptosis in stimulated macrophages

Previous studies demonstrated that nucleotides and nucleic acids are closely involved in the signaling process inducing pyroptosis, a highly inflammatory form of programmed cell death in macrophages that can be triggered by stimulation associated with infection (Nakahira et al., 2011; Shimada et al., 2012; Zhong et al., 2018; Riley and Wg Tait, 2020). Therefore, we next examined the impact of HPRT on pyroptosis. Pyroptosis was induced by treatment with nigericin after wildtype or *Hprt* KO macrophages were primed by LPS+IFNγ. Both *Hprt* KO macrophage lines had greatly increased pyroptosis compared to wildtype; in contrast, in the no priming control, cell death upon sole nigericin treatment remained low and unchanged compared to wildtype cells (Figure 7C). This result suggests that loss of HPRT-dependent purine salvage greatly promotes pyroptosis, suggesting implications of nucleotide metabolism in inflammation driven by macrophages.

#### Purine metabolism impacts T. gondii proliferation in macrophages

Finally, we examined the significance of purine salvage in infection. As one of the first lines of defense against foreign invaders, macrophages are vital for proper clearance of pathogens. However, they can also be hijacked by some pathogens for replication and dissemination. *Toxoplasma gondii* is one type of intracellular parasite that uses infected macrophages as a major means for dissemination across the body (Courret et al., 2006). Interestingly, *T. gondii* are purine auxotrophic; they lack essential purine *de novo* synthesis enzymes and instead acquire the purine nucleotides necessary for their replication by salvaging nucleosides and bases from their host cell (Supplemental figure 7E). Therefore, we hypothesized that altered host purine metabolism can impact *T. gondii* growth in macrophages. To test this hypothesis, we infected wildtype, *Hprt* KO, or *Xor* KO macrophages with the rapidly replicating form of *T. gondii* called a tachyzoite. The growth of *T. gondii* was significantly faster in *Hprt* KO and *Xor* KO macrophages than wildtype (Figure 7D). To further dissect whether the increased pathogen growth in the *Hprt* KO and *Xor* KO are due to more effective initial infection or increased parasite replication rate, we analyzed the number of parasites per vacuole 24h after infection using microscopy. Because the initial infection results in one parasite per vacuole in host macrophages, the number of vacuoles containing two, four, or eight parasites indicates *T. gondii* replication in the vacuole. We observed more vacuoles containing two or more parasites after 24h in *Xor* KO macrophages, and to a greater extent in *Hprt* KO macrophages (Figure 7E). This result shows that *T. gondii* replicates faster when host *Hprt* or *Xor* is lost, likely because the activities of host HPRT and XOR can compete with the parasites for purine substrates, ultimately limiting parasite growth. It demonstrated that beyond regulating host functions, the rewiring of macrophage nucleotide metabolism can allow cells to reroute key metabolites that could otherwise be used by pathogens, impacting host–pathogen interactions.

## Discussion

This study revealed that upon stimulation with classical activation signals, macrophages undergo substantial, systematic reprogramming of nucleotide metabolism. Specifically, *de novo* synthesis of purine and most pyrimidine nucleotides is shut down, nucleotide degradation is increased, and cells switch to salvage to maintain most nucleotides. Unlike in T-cells, where the remodeling of nucleotide metabolism has been highlighted and found largely related to the increased metabolic demands associated with their rapid expansion upon immune activation (Toy et al., 2010; Sauer et al., 2012; Ron-Harel et al., 2016; Vigano et al., 2019), primary macrophages (whether activated or not) do not proliferate, and thus have no net nucleotide requirement for biomass expansion. Nonetheless, our data showed general nucleotide turnover is active in both stimulated and unstimulated macrophages, and rewiring of nucleotide metabolism upon stimulation is functionally significant. So, a question from the flux balance perspective is: why do non-proliferating immune cells need dynamic nucleotide metabolism?

One answer lies in the close connection between nucleotide metabolism and many other metabolic pathways, and their involvement in a range of cellular processes. For instance, nucleotides are crucial for supporting transcription, and transcriptional reprogramming is highly active upon immune activation; nucleotides are precursors of essential cofactors such as SAM and NAD, whose metabolic rewiring is important for macrophage immune response and function (Yu et al., 2019; Minhas et al., 2019; Cameron et al., 2019); and many nucleotides and nucleosides, especially purines such as ATP, adenosine, and inosine, are released into the extracellular space and act as immunoregulatory signals (Linden et al., 2019; Antonioli et al., 2018; Haskó et al., 2004). Therefore, the changes in purine nucleotide metabolism, particularly the release and uptake of extracellular purines, can have significant impacts on autocrine and paracrine signaling.

Second, although non-proliferating cells do not have a net demand for DNA for growth, mitochondria replication and turnover are still active, which require dNTPs. Mitochondrial DNA has been found to be important for inflammasome activation in macrophages (Nakahira et al., 2011; Shimada et al., 2012; Zhong et al., 2018; Riley and Wg Tait, 2020). Interestingly, we found thymidine nucleotides, the only nucleotide type needed for DNA but not for RNA, is uniquely depleted. It is possible the nucleotide imbalance has important effects in intracellular signaling, and in mitochondrial DNA synthesis or fragmentation. Moreover, we have also made an intriguing observation from transcriptomic data that changes in nucleotide metabolism genes appear to be compartmentalized— some mitochondrial nucleotide metabolism genes, such as *Cmpk2* and *Ak4*, are strongly upregulated upon stimulation, while their cytosolic counterparts are not (Figure 1E). Mitochondrial versus cytosolic nucleotide metabolism can have different downstream impacts on nucleic acid metabolism and other processes. The questions regarding compartmentalized nucleotide balance merit more research in the future.

Finally, macrophages perform essential functions in pathogen elimination, wound repair, and inflammation regulation, and nucleotide metabolism is involved in macrophages’ interactions with the microenvironment. During infection, macrophages can phagocytose and digest pathogens. In this case, increased nucleotide degradation capacity could prepare macrophages for the task, and the switch from nucleotide *de novo* synthesis to salvage can be beneficial to avoid nucleotide overload. Macrophages are also important for clearing apoptotic cells and neutrophil extracellular traps, which are a network of cellular DNA mixed with anti-bacterial proteins released by neutrophils as a means of anti-infection defense. Macrophages’ ability to effectively clear these nucleic acid-rich host components is critical for tissue homeostasis and inflammation control. Indeed, it has been recently reported that classically activated macrophages have greater capability to clear neutrophil extracellular traps than unstimulated macrophages (Haider et al., 2020), which is likely coupled to changes in nucleotide metabolism, such as the increased nucleotide degradation and salvage capacity we revealed here. Moreover, emerging research showed that in tumor microenvironments, changes in macrophage metabolism can impact cancer cells’ survival in response to a chemotherapy drug that targets nucleotide metabolism (Halbrook et al., 2019), exemplifying the importance of nucleotide metabolism in macrophage–cancer interactions. Thus, the stimulation-induced reprogramming of nucleotide metabolism can have effects ranging from the support and regulation of cell-intrinsic functions to metabolic interactions between macrophages and pathogens, other immune cells, and cancer cells.

Our study also pinned down the key regulation points driving the rewiring of nucleotide metabolism upon classical stimulation, and identified the mechanisms by which they are regulated. Specifically, the shutdown of purine nucleotide *de novo* synthesis is driven by a combination of transcriptional downregulation of *de novo* synthesis genes (minor contribution) and NO-driven inhibition of the key enzyme ATIC (major rate-limiting factor). Interestingly, we found the transcription of many nucleotide synthesis genes are down regulated in a temporally correlated manner. It is likely a master regulator coordinates changes in these genes. One possible candidate is the transcription factor *Myc*, which is known to positively regulate many nucleotide synthesis genes (Lane and Fan, 2015). Here, we found its expression is significantly reduced soon after LPS+IFNγ stimulation (Supplemental figure 8A), correlating with its target genes in nucleotide *de novo* synthesis. The coordinated transcriptional downregulation could also be beneficial when the last enzyme in purine *de novo* synthesis, ATIC, is profoundly inhibited, thus limiting overall purine synthesis flux in stimulated macrophages. In this case, the transcriptional downregulation of upstream genes can prevent continued overproduction of purine synthesis intermediates, as supported by the increase-then-subside dynamics of AICAR we observed here (Supplemental figure 6D).

To our knowledge, the profound inhibition of ATIC by NO has not been reported in literature before, and the specific molecular mechanism is not yet clear. However, we found that manipulating cellular NO level by *Nos2* knockout did not have a major impact on ATIC total protein level (Supplemental figure 8B), even though it significantly restored ATIC flux in stimulated macrophages (Figures 6C–D). Together these observations suggest that the mechanism is not mainly through NO altering the synthesis of the ATIC enzyme or promoting ATIC degradation. We hypothesize the most likely mechanism is through a post-translational modification. In the catalytic center of the ATIC enzyme, there is a cysteine residue (C101) which is critical for FAICAR binding and catalyzing IMP cyclohydrolase activity (Szabados et al., 1994; Uniprot). Using the computational prediction algorithm DeepNitro (Xie et al., 2018), we found this cysteine residue is predicted to be one of the most susceptible to nitrosylation by NO (Supplemental figure 8C). Indeed, a recent proteomic analysis in human induced pluripotent stem cell-derived neurons reported this site of ATIC can be nitrosylated (Doulias et al., 2023). It is likely such nitrosylation drives the inhibition of ATIC when NO production is induced upon macrophage activation. Similarly, we found the enzyme XOR is also profoundly inhibited by NO. The catalytic mechanism of XOR requires two Fe-S clusters to transfer electrons (Berry and Hare, 2003). It is possible that NO can disrupt the Fe-S clusters, thereby shutting down complete purine degradation activity of XOR, which, as we showed above, is critical for diverting flux to support salvage instead. Further work is merited to test these hypotheses.

Here we showed that purine nucleotide salvage has significant impacts in macrophage migration, pyroptosis, and *T. gondii* replication in macrophages, demonstrating the functional importance of macrophage nucleotide metabolism. Such significance and health relevance are likely broader than what has been directly assessed here. For instance, *T. gondii* just presents one example of an intracellular protozoan parasite that is a purine auxotroph, but there are multiple others, including the pathogens causing diseases such as leishmaniasis and malaria. This raises an important point regarding macrophages’ role in infections. As one of the first lines of defense against foreign invaders, macrophages are vital for proper clearance of pathogens. However, they can also be hijacked by some pathogens for replication and dissemination. It is therefore crucial to evaluate the interaction between macrophage and pathogen metabolisms in specific and diverse contexts.

The impact of nucleotide metabolism on macrophage pyroptosis and migration also has broad implications. Pyroptosis is a part of the antimicrobial response, but due to its highly inflammatory nature, pyroptosis can also cause tissue damage and is associated with many diseases (Wei et al., 2021). Interestingly, we found that HPRT activity appears to help control pyroptosis, and the flux through HPRT is increased in macrophages after prolonged stimulation, when cells are transitioning to a less inflammatory state. This points to the possibility that switching to increased purine salvage may help control inflammation and promote resolution. The exact molecular mechanism connecting purine salvage and pyroptosis control, and its implications in inflammatory conditions, merits future studies. We hypothesize a possible mechanism is through the activation of AMPK. Among the many functions of AMPK signaling is the inhibition of the inflammasome (Cordero et al., 2018; Yang et al., 2019), which is required for pyroptosis. We found AMPK is increasingly phosphorylated and thus strongly activated upon macrophage stimulation, and such activation is significantly reduced in stimulated *Hprt* KO cells (Supplemental figure 8D). *Hprt* KO may therefore de-repress pyroptosis. However, more in-depth testing of this hypothesis is required, although beyond the scope of this study.

Overall, this study thoroughly revealed the substantial rewiring of nucleotide metabolism in macrophages upon classical activation, identified its underlying mechanisms, and illustrated its functional significance. It further maps out avenues for many future discoveries in both the realms of molecular regulation mechanisms and of clinical relevance and translation.

## Figure Legends

**Supplemental figure 1:**
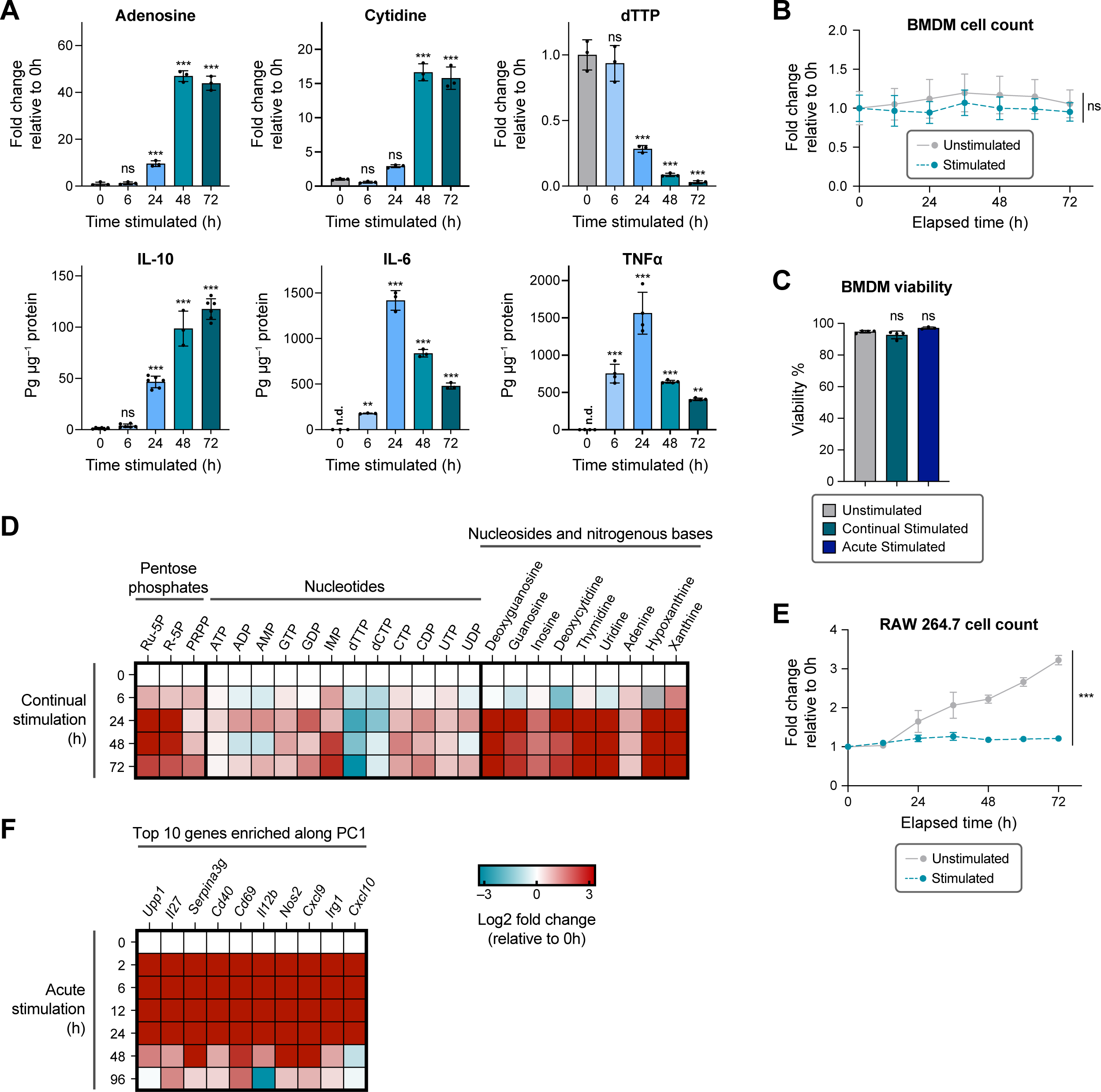
A) Cooccurrence between the shift in nucleotide metabolism and cytokine production. Relative intracellular abundance of adenosine, cytidine, and dTTP are shown as examples of the dynamic changes in nucleotide metabolism in BMDMs over a timecourse of continual stimulation, with full changes shown in Figure 1C. Mean ± SD (n=3 independent samples for each). *The dynamic changes in IL-6 and TNFα production are re-represented from data reported in our previous publication (Seim et al., 2019) to show the correlation. B) Relative cell number of BMDMs over a timecourse with or without continual stimulation. Mean ± SD (n=4 independent samples). C) Cell viability, as determined by the percentage of cells negative for Cytotox green dye, in a population of BMDMs that are unstimulated and cultured for 72h, continually stimulated for 72h, or acutely stimulated for 96h. Mean ± SD (n=4 independent samples for unstimulated and continually stimulated; n=3 independent samples for acute stimulated). D) Metabolomic changes in pentose phosphates, nucleotides, nucleosides, and nitrogenous bases in RAW 264.7 cells over a timecourse of continual stimulation. Relative metabolite abundances are compared to unstimulated cells (0h) and displayed on a log2 scale as a heatmap. Each box represents the mean of n=3 independent samples. E) Relative cell number of RAW 246.7 cells over a timecourse with or without continual stimulation. Mean ± SD (n=3 independent samples). F) The top 10 genes enriched along principal component 1 of the transcriptomic changes in BMDMs over a timecourse of acute stimulation. The relative expression levels are compared to unstimulated cells (0h) and displayed on a log2 scale as a heatmap. Each box represents the mean of n=3 independent samples. B, E) Statistical analysis was performed with unpaired student’s t test comparing unstimulated to stimulated conditions at 72h. A, C) Statistical analysis was performed using one-way ANOVA followed by post hoc Dunnett’s test comparing all groups to unstimulated cells. ** indicates p<0.01; *** indicates p<0.001; ns indicates not significant (p>0.01).

**Supplemental figure 2:**
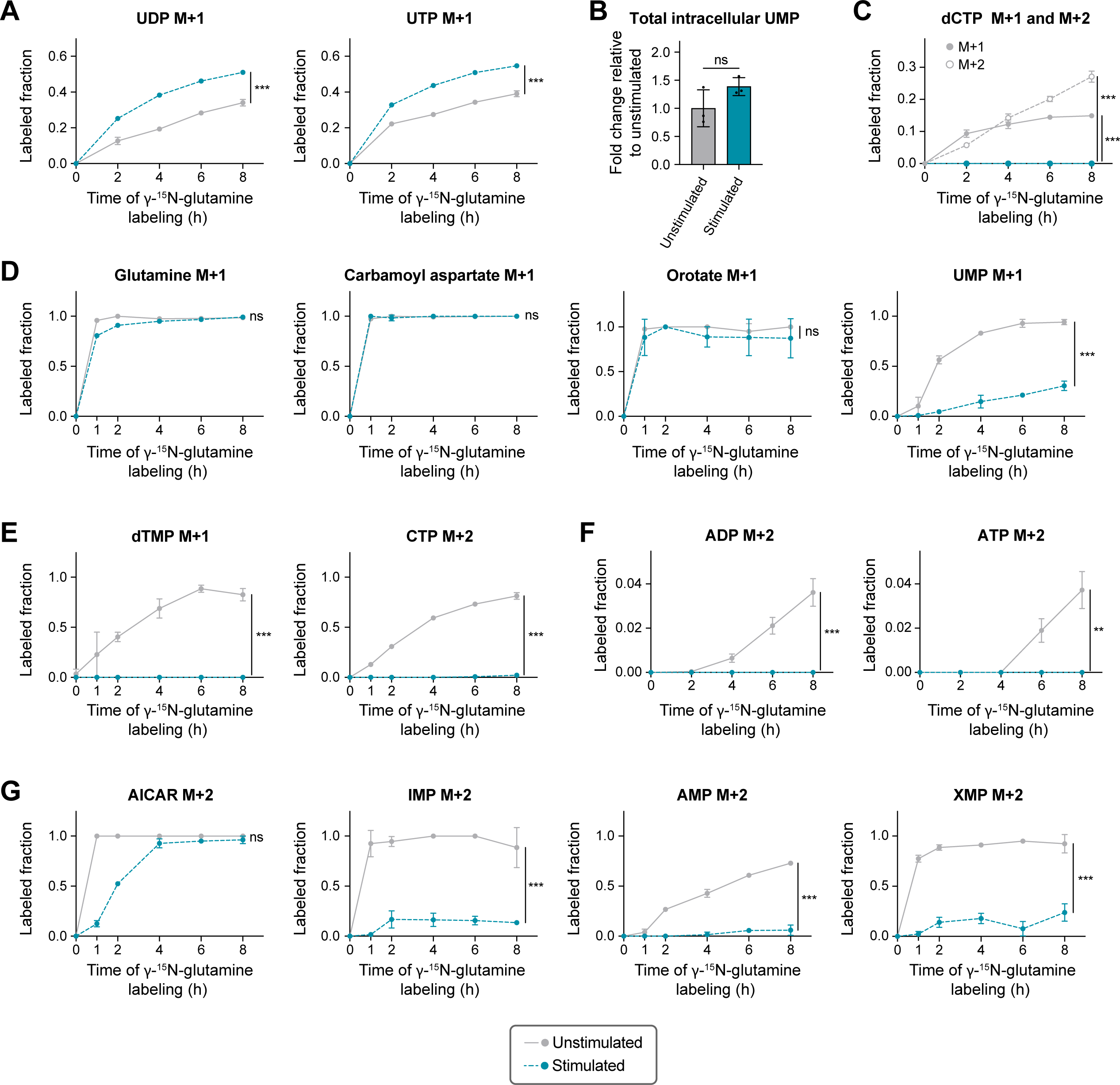
A) Labeling kinetics of UDP and UTP in unstimulated or stimulated BMDMs. B) Relative intracellular abundance of UMP in unstimulated or stimulated BMDMs. Mean ± SD (n=3 independent samples). Data are presented relative to abundance in unstimulated cells. C) Labeling kinetics of dCTP in unstimulated or stimulated BMDMs. D–E) Labeling kinetics of (D) glutamine, carbamoyl aspartate, orotate, UMP and (E) dTMP, CTP in unstimulated or stimulated RAW 264.7 macrophages. F) Labeling kinetics of ADP and ATP in unstimulated or stimulated BMDMs. G) Labeling kinetics of AICAR, IMP, AMP, and XMP in unstimulated or stimulated RAW 264.7 macrophages. A, C–G) Cells were labeled with γ-^15^N-glutamine for various time points as indicated on x-axis. Mean ± SD (n=3 independent samples). A–C, F) Stimulated cells are continually stimulated for 48h. D–E, G) Stimulated cells are continually stimulated for 24h. A-G) Statistical analysis was performed with unpaired student’s t test comparing unstimulated to stimulated cells at the final time point. ** indicates p<0.01; *** indicates p<0.001; ns indicates not significant (p>0.01).

**Supplemental figure 3:**
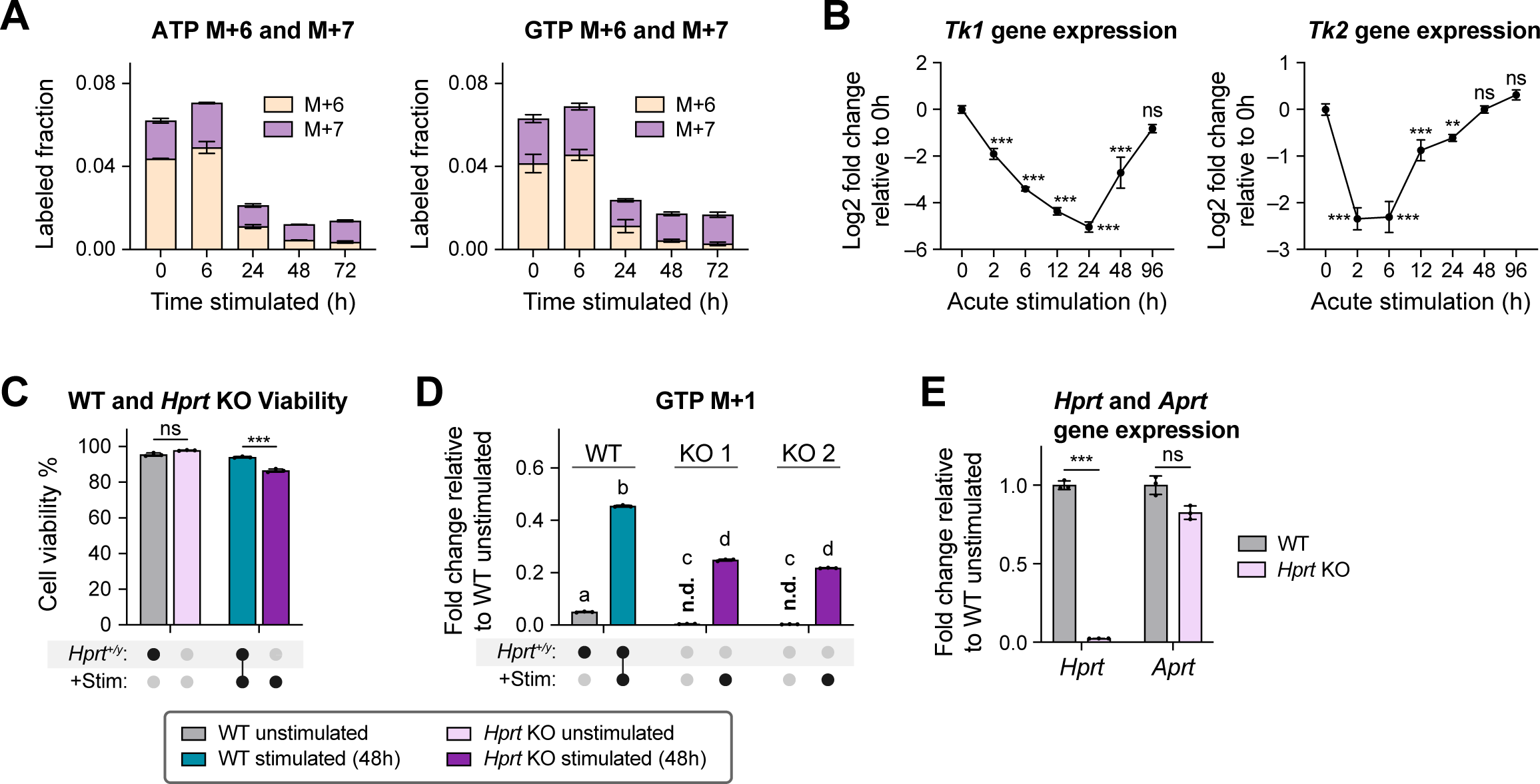
A) Fractions of M+6 and M+7 labeled ATP and GTP in BMDMs over a timecourse of continual stimulation. Cells were labeled with U-^13^C-glucose for 24h before analysis. B) Transcriptional changes of thymidine kinase *Tk1* and *Tk*2 in BMDMs over a timecourse of acute stimulation. Relative expression is compared to unstimulated cells (0h) and displayed on a log2 scale. C) Cell viability, as determined by the percentage of cells negative for Cytotox green dye, in wildtype and *Hprt* KO RAW 264.7 cells, either unstimulated or continually stimulated for 48h. D) M+1 labeled fraction of GTP in wildtype or *Hprt* KO (2 separate clones) RAW 264.7 cells, either unstimulated or continually stimulated for 48h. Cells were labeled with γ-^15^N-glutamine for 24h before analysis. E) Expression levels of *Hprt* and *Aprt* (encoding adenine phosphoribosyltransferase) in unstimulated wildtype or *Hprt* KO RAW 264.7 cells. Relative expression of each gene is compared to wildtype. A–E) Mean ± SD (n=3 independent samples). C, E) Statistical analysis was performed with unpaired student’s t test comparing wildtype to *Hprt* KO. B) Statistical analysis was performed using one-way ANOVA followed by post hoc Dunnett’s test comparing all groups to unstimulated cells (0h). ** indicates p<0.01; *** indicates p<0.001; ns indicates not significant (p>0.01). D) Statistical analysis was performed using one-way ANOVA followed by post hoc Tukey’s test. Bars with different lower-case letters (a, b, c, or d) indicate a statistically significant difference with p<0.05.

**Supplemental figure 4:**
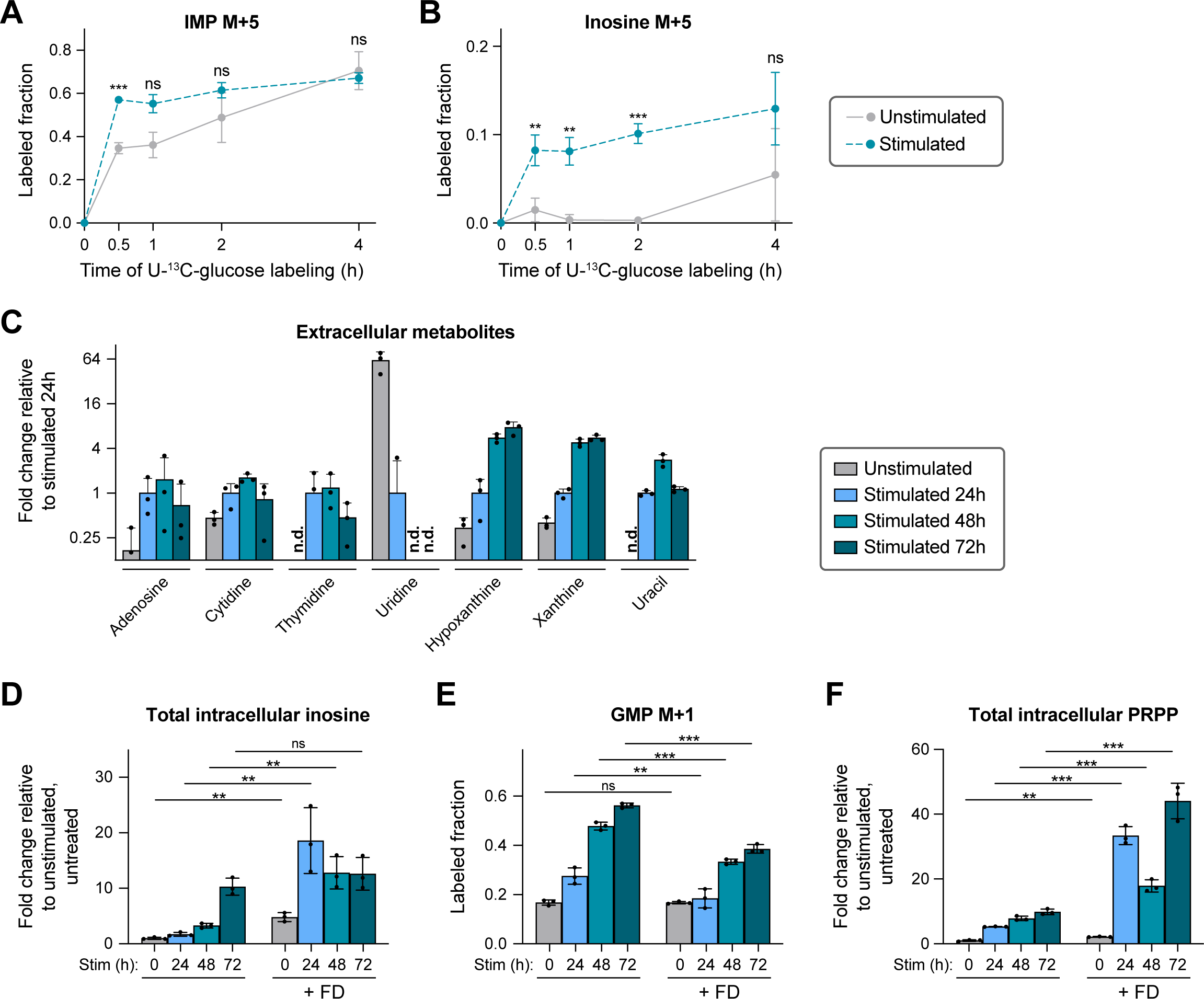
A–B) Labeling kinetics of (A) IMP and (B) inosine in unstimulated or 48h continually stimulated BMDMs after labeling with U-^13^C-glucose for 0–4 hours. C) Extracellular metabolites released into the media by BMDMs over a timecourse of continual stimulation. Cells were incubated in the media for 22h before analysis. Data are presented relative to the level in spent media incubated with 24h stimulated BMDMs on a log scale. N.d. indicates not detected. Mean + SD (n=3 independent samples). D) Relative intracellular abundance of inosine in BMDMs over a timecourse of continual stimulation, with or without treatment of 10 µM forodesine (FD) throughout the timecourse. Data are presented relative to abundance in unstimulated, untreated cells. E) Fraction of M+1 labeled GMP from 24h γ-^15^N-glutamine labeling in BMDMs over a timecourse of continual stimulation, with or without treatment of 10 µM forodesine (FD) throughout the timecourse. F) Relative intracellular PRPP abundance in BMDMs over a timecourse of continual stimulation, with or without treatment of 10 µM forodesine (FD) throughout the timecourse. Data are presented relative to abundance in unstimulated, untreated cells. A–B, D–F) Mean ± SD (n=3 independent samples). A–B) Statistical analysis was performed with unpaired student’s t test comparing unstimulated to stimulated cells at each time point. D–F) Statistical analysis was performed with unpaired student’s t test comparing untreated to FD treated cells at each time point. ** indicates p<0.01; *** indicates p<0.001; ns indicates not significant (p>0.01).

**Supplemental figure 5:**
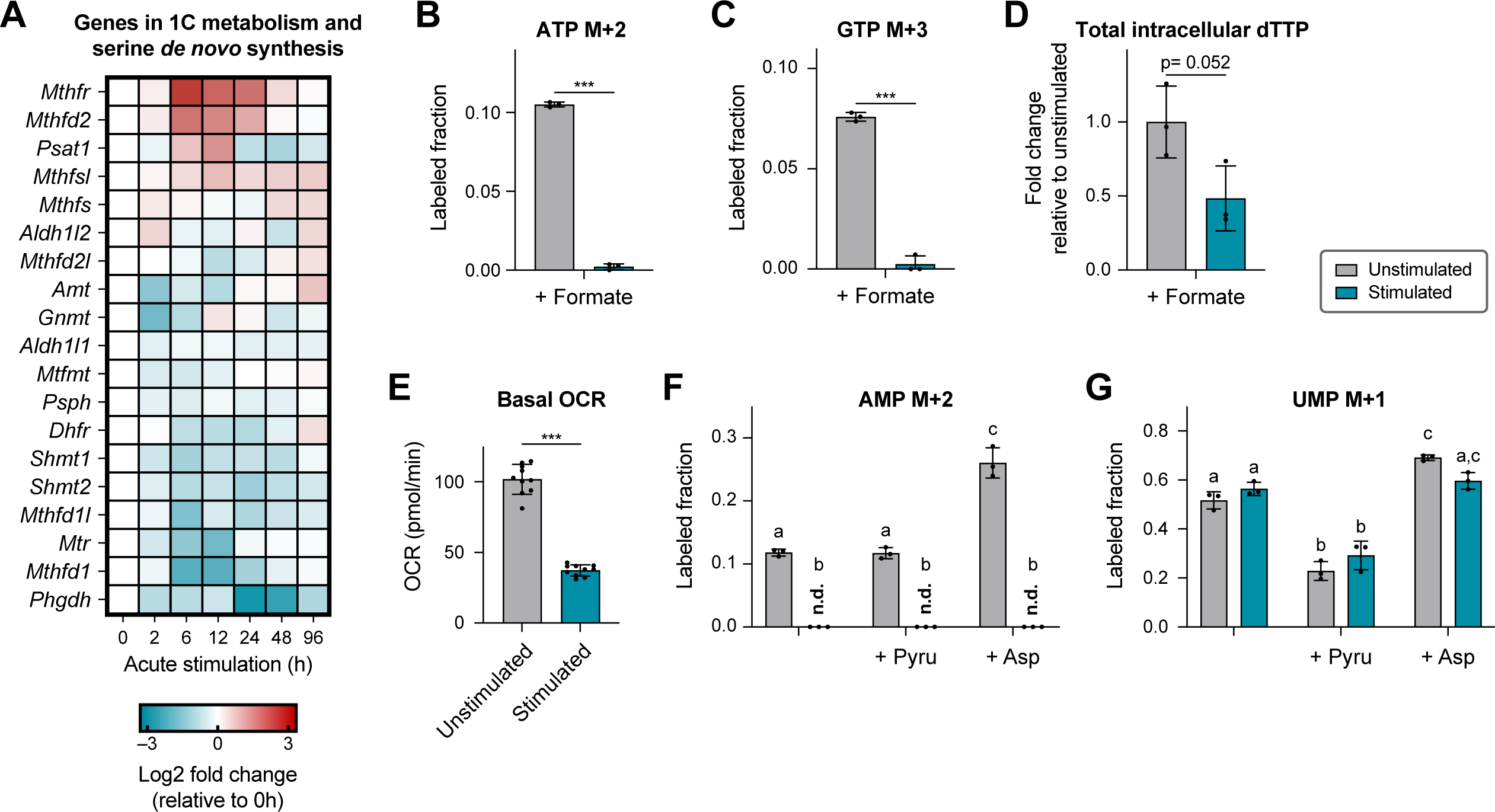
A) Transcriptional changes of detected genes in one carbon metabolism and serine *de novo* synthesis in BMDMs over a timecourse of acute stimulation. Relative expression of each gene is presented as log2 fold change compared to unstimulated cells (0h) and displayed as a heatmap. Each box represents the mean of n=3 independent samples. B–C) Labeled fraction of (B) M+2 labeled ATP and (C) M+3 labeled GTP in unstimulated or stimulated BMDMs supplemented with 1mM ^13^C-formate for 24h. D) Relative total intracellular dTTP abundance in unstimulated or stimulated BMDMs supplemented with 1mM ^13^C-formate for 24h. Data are presented relative to abundance in unstimulated cells. E) Basal oxygen consumption rate of unstimulated or stimulated BMDMs. Mean ± SD (n=10 independent samples). F–G) Labeled fraction of (F) M+2 labeled AMP and (G) M+1 labeled UMP after 24h γ-^15^N-glutamine labeling in unstimulated or stimulated BMDMs, with or without supplementation of 2.5mM pyruvate or 5mM aspartate throughout the timecourse. N.d. indicates not detected. B–G) Stimulated cells are continually stimulated for 48h. B–D, F–G) Mean ± SD (n=3 independent samples). B–E) Statistical analysis was performed with unpaired student’s t test comparing unstimulated to stimulated cells. *** indicates p<0.001. F–G) Statistical analysis was performed using one-way ANOVA followed by post hoc Tukey’s test. Bars with different lower-case letters (a, b, or c) indicate a statistically significant difference with p<0.05.

**Supplemental figure 6:**
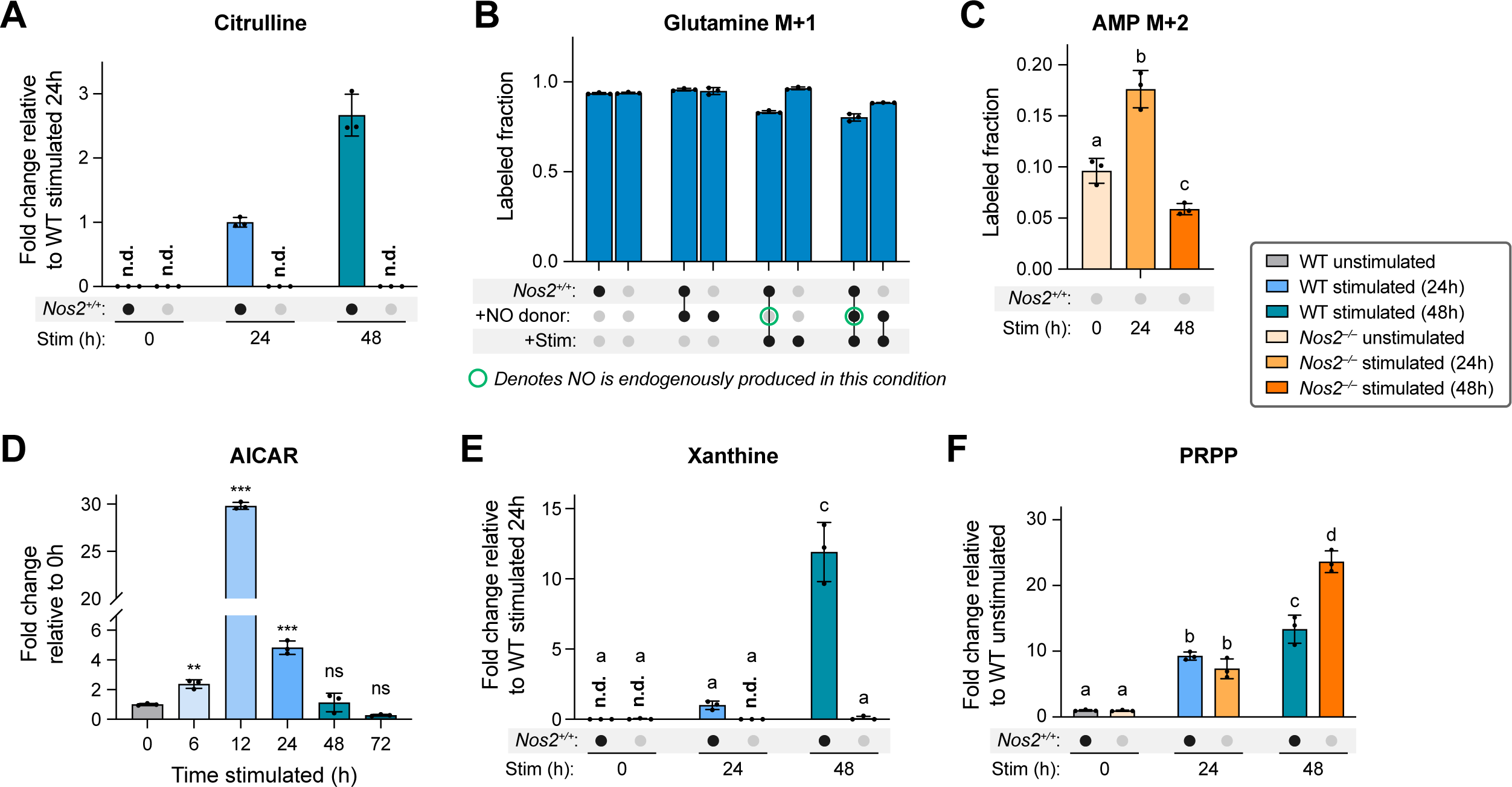
A) Relative intracellular abundance of citrulline in wildtype or *Nos2*^-/-^ BMDMs over a timecourse of continual stimulation. Data are presented relative to abundance in 24h stimulated wildtype cells. N.d. indicates not detected. B) Fraction of M+1 labeled glutamine from 24h γ-^15^N-glutamine labeling in unstimulated or 48h continually stimulated wildtype or *Nos2*^-/-^ BMDMs, with or without treatment of 200 µM NO donor DETA-NONOate throughout the timecourse. C) Fraction of M+2 labeled AMP from 24h γ-^15^N-glutamine labeling in *Nos2*^-/-^ BMDMs over a timecourse of continual stimulation. D) Relative intracellular AICAR abundance in BMDMs over a timecourse of continual stimulation. Data are presented relative to abundance in unstimulated cells (0h). E–F) Relative intracellular abundance of (E) xanthine and (F) PRPP in wildtype or *Nos2*^-/-^ BMDMs in (E) WT 24h stimulated cells and (F) WT unstimulated cells. N.d. indicates not detected. A–F) Mean ± SD (n=3 independent samples). D) Statistical analysis was performed using one-way ANOVA followed by post hoc Dunnett’s test comparing all groups to unstimulated cells (0h). ** indicates p<0.01; *** indicates p<0.001; ns indicates not significant (p>0.01). C, E–F) Statistical analysis was performed using one-way ANOVA followed by post hoc Tukey’s test. Bars with different lower-case letters (a, b, c, or d) indicate a statistically significant difference with p<0.05.

**Supplemental figure 7:**
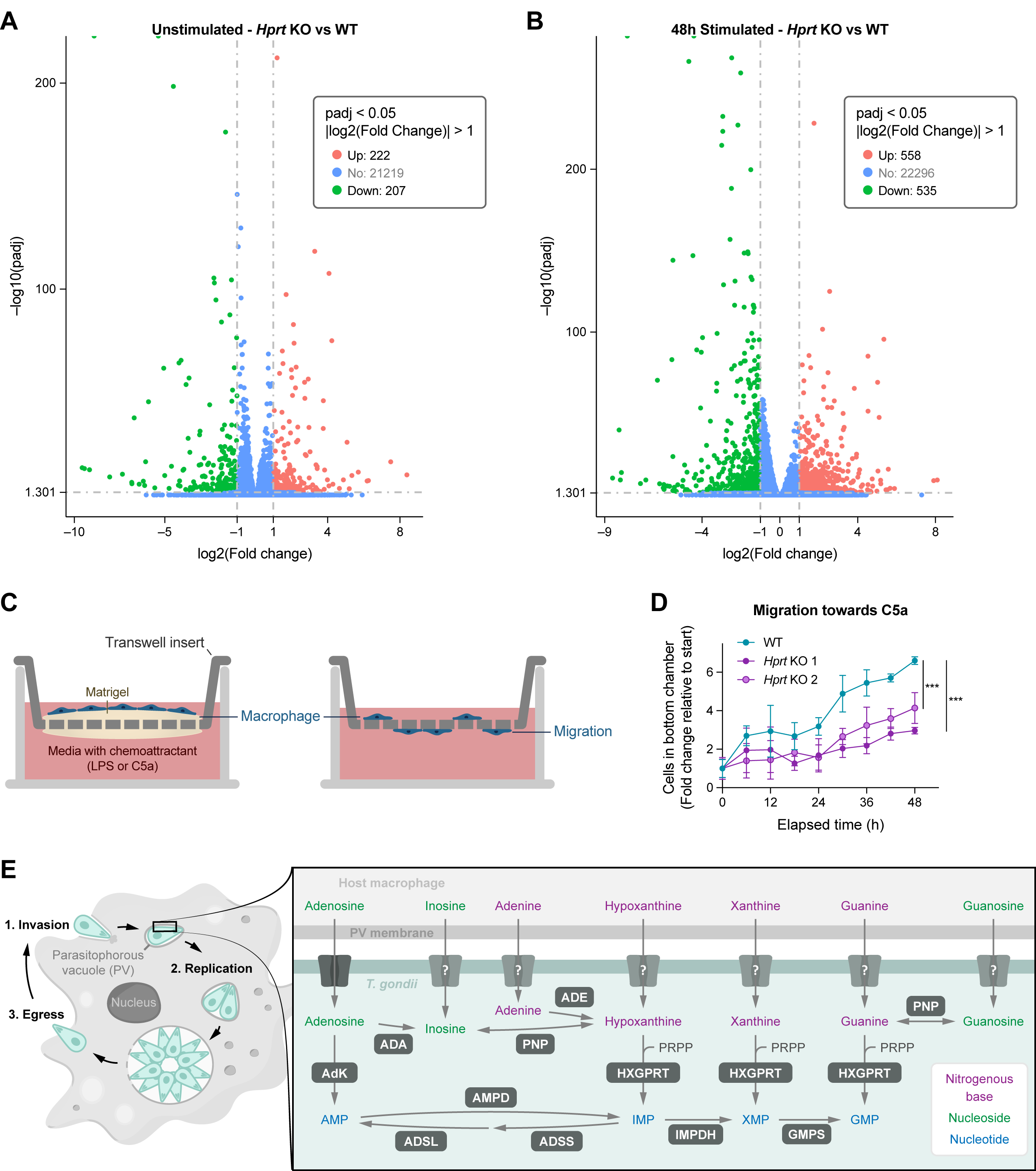
A–B) Volcano plot showing differentially expressed genes (up-regulated in red and down-regulated in green) in *Hprt* KO RAW 264.7 cells compared to wildtype cells in A) unstimulated state, and B) 48h stimulated state. C) Schematic of a macrophage migration assay. After a chemoattractant is added, macrophages break down the Matrigel and migrate to the other side of the transwell. D) Migration of wildtype and *Hprt* KO (2 clones) RAW 264.7 cells across a Matrigel-coated transwell towards C5a. Mean ± SD (n=5 independent samples). Statistical analysis was performed using one-way ANOVA followed by post hoc Dunnett’s test comparing *Hprt* KO to wildtype. *** indicates p<0.001. E) Schematic showing *T. gondii* replication cycle in a macrophage and its dependence on salvaging host purine nucleosides and bases. Specifically identified is the parasitophorous vacuole that separates host cell cytosol from *T. gondii*.

**Supplemental figure 8:**
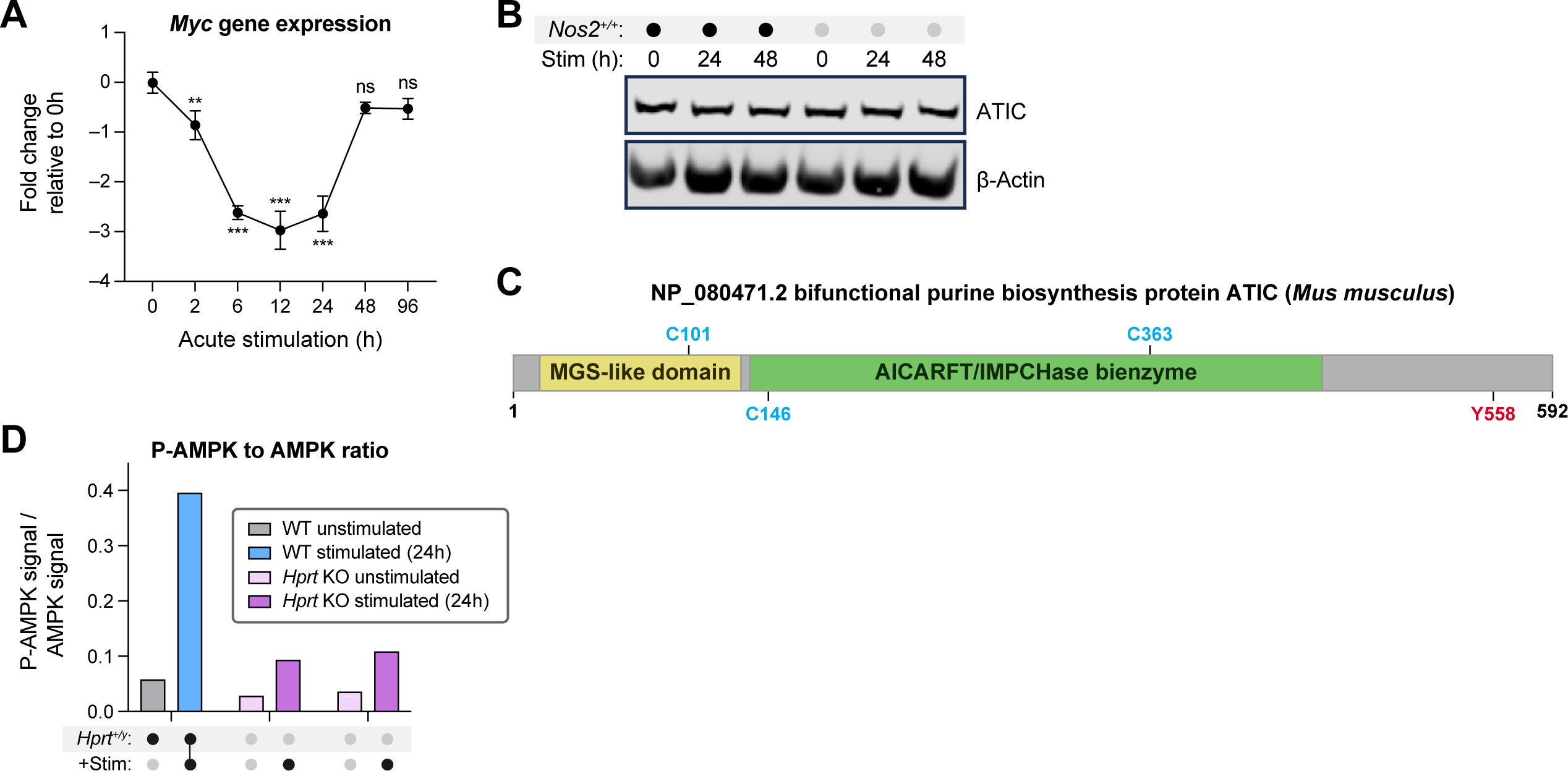
A) *Myc* expression in BMDMs over a timecourse of acute stimulation. Data are presented relative to expression in unstimulated cells (0h) and displayed on a log2 scale. Mean ± SD (n=3 independent samples). Statistical analysis was performed using one-way ANOVA followed by post hoc Dunnett’s test comparing all groups to unstimulated cells (0h). ** indicates p<0.01; *** indicates p<0.001; ns indicates not significant (p>0.01). B) ATIC protein levels in wildtype or *Nos2*^-/-^ BMDMs over a timecourse of continual stimulation. C) Predicted ATIC nitrosylation sites by DeepNitro. D) The ratio between P-AMPK (Thr 172) to AMPK protein level, quantified from immunoblotting in unstimulated or 24h continually stimulated wildtype or *Hprt* KO (2 separate clones) RAW 264.7 cells.

## Methods

### Cell Culture

Murine bone marrow-derived macrophages (BMDMs) were isolated from wildtype C57BL/6J mice or B6.129P2-Nos2tm1Lau/J (*Nos2*^-/-^) mice (Jackson Lab), as indicated. Mice were bred and maintained according to protocols approved by the University of Wisconsin–Madison Institutional Animal Care and Use Committee. Bone marrow cells were isolated and differentiated as previously described (Seim et al., 2020) from 6–12-week-old males or females. BMDMs were cultured in RPMI 1640 (VWR: 16750-084) with 10% dialyzed fetal bovine serum (VWR: 16777-212), 25mM HEPES, 2mM glutamine, 1% antibiotics (Thermo Fisher Scientific: 15140122-100 IU/mL penicillin and 100ug/mL streptomycin) and 20ng/mL M-CSF (R&D Systems: 416-ML) at 37°C and 5% CO_2_. To stimulate BMDMs, cells were incubated with 50ng/mL LPS (E. coli O111:B4, Sigma: L3024) and 30ng/mL recombinant mouse IFNγ (R&D Systems: 485-MI).

Wildtype RAW 264.7 cells (ATCC) or knockouts were cultured in RPMI 1640 with 10% dialyzed fetal bovine serum, 25mM HEPES, 2mM glutamine, 1% antibiotics at 37°C and 5% CO_2_. To stimulate the cells, 50ng/mL LPS and 10ng/mL recombinant mouse IFNγ were added to the media.

During cell culture experiments for both RAW 264.7 and BMDMs, media was refreshed daily to avoid the depletion of nutrients and to maintain LPS+IFNγ stimulation. Media was also refreshed 2–4h before collections. For time course experiments, the time of starting stimulation was staggered to allow all treatment groups to be collected at the same time and control for the potential impact of culturing cells for different durations. For continual stimulation experiments, LPS and IFNγ were maintained in the media throughout. For acute stimulation experiments, cells were cultured with LPS and IFNγ for 2h, after which stimuli were removed, cells were washed with DPBS, and cell culture was continued in fresh media without stimuli for the remaining timecourse.

Cell number and viability were measured using two assays. When plating cells for experiments, RAW 264.7 and BMDMs were detached using Accutase solution (Sigma-Aldrich: A6964), pelleted and resuspended in media. Then an aliquot of the cell suspension was taken and stained with an equal volume of Trypan Blue (Gibco: 15250-061), and live/dead cell number was assessed using a Countess II Automated Cell Counter (Thermo Fisher). To measure cytotoxicity in live culture, 20–40,000 cells were plated per well of a 96-well plate, stained with 250nM Cytotox green (Sartorius: 4633), and imaged using an Incucyte S3 Live-Cell Analysis System (Sartorius).

### Cytokine release assay

For cytokine release measurements, 22h spent media from cells was spun at 500xg for 5 minutes and the supernatant was collected and frozen until further analysis. IL-10, TNF-α and IL-6 release was measured using DuoSet ELISA kits according to the manufacturer’s instructions (R&D Systems: DY417-05; DY410-05; DY406-05).

### Metabolomics analysis

To measure intracellular metabolites, cells were washed twice with DPBS and metabolites were extracted with -80°C LC-MS grade 80:20 MeOH:H_2_O (Fisher Scientific: A456-4; W64). The extract was transferred to a tube, vortexed, and centrifuged at max speed. Then the supernatant was transferred to a new tube. The remaining pellet was extracted again with 80:20 MeOH:H_2_O and both supernatants were combined and dried under nitrogen stream. Dried samples were resuspended in LC-MS grade H_2_O or LC-MS grade 40:40:20 ACN:MeOH:H_2_O (ACN-Fisher Scientific: A955-4) as loading buffer depending on the LC-MS method.

To measure extracellular metabolites, spent media samples were extracted with 4x volume of LC-MS grade MeOH and vortexed. The extract was centrifuged at max speed. The supernatant was further diluted 1:10 with either LC-MS grade H_2_O or LC-MS grade 40:40:20 ACN:MeOH:H_2_O depending on the LC-MS method.

Samples were analyzed using a Thermo Q-Exactive mass spectrometer coupled to a Vanquish Horizon UHPLC. Two methods were used for optimal quantification.

In the first method, samples resuspended in LC-MS grade H_2_O were separated on a 100 × 2.1 mm, 1.7μM Acquity UPLC BEH C18 Column (Waters) with a gradient of solvent A (97:3 H_2_O:MeOH, 10mM TBA (Sigma Aldrich: 90781), 9mM acetic acid (Fisher Scientific: A11350), pH 8.2) and solvent B (100% MeOH). The gradient is: 0 min, 5% B; 2.5 min, 5% B; 17 min, 95% B; 21 min, 95% B; 21.5 min, 5% B. The flow rate is 0.2mL/min with a 30°C column temperature. Data were collected on a full scan negative mode at a resolution of 70K. Settings for the ion source were: 10 aux gas flow rate, 35 sheath gas flow rate, 2 sweep gas flow rate, 3.2kV spray voltage, 320°C capillary temperature and 300°C heater temperature.

In the second method, samples resuspended in LC-MS grade 40:40:20 ACN:MeOH:H_2_O were separated on a 2.1 × 150mm Xbridge BEH Amide (2.5 μM) Column (Waters) using a gradient of solvent A (95% H_2_O, 5% ACN, 20 mM ammonium acetate, 20 mM NH_4_OH) and solvent B (20% H_2_O, 80% ACN, 20 mM ammonium acetate, 20 mM NH_4_OH). The gradient is: 0 min, 100% B; 3 min, 100% B; 3.2 min, 90% B; 6.2 min, 90% B; 6.5 min, 80% B; 10.5 min, 80% B; 10.7 min, 70% B; 13.5 min, 70% B; 13.7 min, 45% B; 16 min, 45% B; 16.5 min, 100% B; 22 min, 100% B. The flow rate is 0.3mL/min with a 30°C column temperature. Data were collected on a full scan positive or negative mode.

Metabolites reported were identified based on exact *m/z* and retention times determined with standards. Data were analyzed with MAVEN (Melamud et al., 2010; Clasquin et al., 2012). Metabolite levels were normalized to either protein content or cell counts. To quantify the release of extracellular metabolites, absolute concentrations were determined using calibration curves generated with standards of the corresponding metabolites.

### Metabolic pathway enrichment analysis

Pathway enrichment analysis was performed using MetaboAnalyst (Xia et al., 2009). Normalized metabolomics data from unstimulated and stimulated (continually stimulated with LPS+IFNγ for 24h) BMDMs were inputted into pathway analysis, log transformed, enriched using global test, and analyzed with the out-degree centrality topology.

### Expression analysis by qPCR

For RNA collection, cells were washed twice with DPBS and then lysed with RNA STAT 60 (Fisher Scientific: NC9884083). Chloroform (Sigma-Aldrich: 650498) was used for phase separation. RNA was precipitated with isopropanol (Fisher Scientific: AC327272500), washed with 75% ethanol (VWR: 71006-012), and solubilized in DEPC H_2_O (VWR: 101076-146). RNA samples were treated with RQ1 DNAse (Promega: M6101) and cDNA was synthesized using SuperScript III Reverse Transcriptase (Thermo Fisher Scientific: 18080093) according to the manufacturer’s instructions. Quantitative PCR was performed on a Roche LightCycler 480 using KAPA SYBR FAST qPCR MasterMix (Roche: KK4611).

Relative gene expression level was normalized to CycloB. The following primers (IDT) were used:

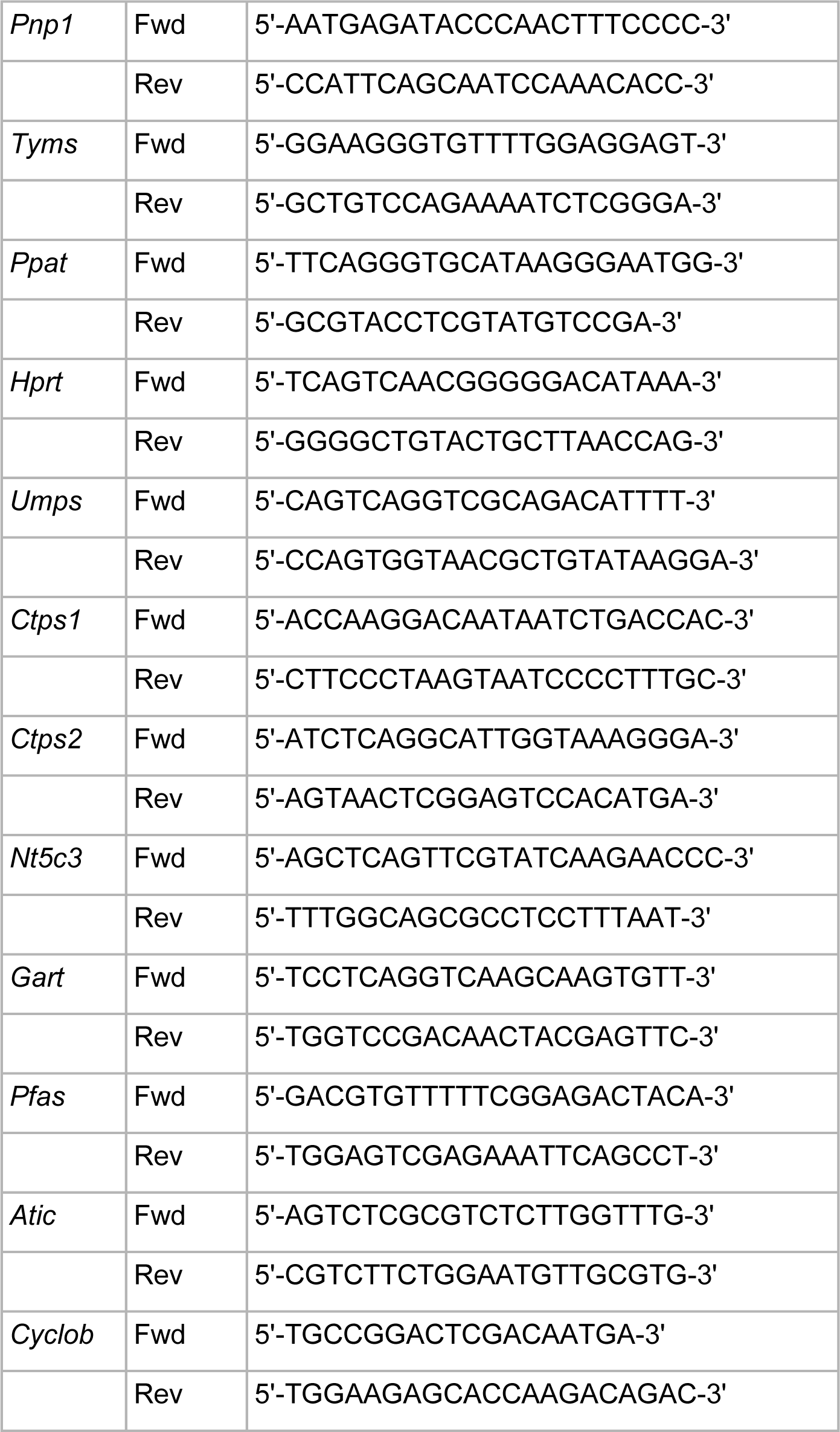

### RNAseq Analysis

RNA sequencing was performed by Novogene, utilizing Illumina’s Sequencing by Synthesis (SBS) technology. RNA samples were collected from BMDMs and RAW 264.7 cells by the same collection method used for qPCR analysis, then submitted for sequencing analysis. Quality control was commenced with base recognition using CASAVA, followed by error rate assessment and GC content analysis. The data were then subjected to stringent filtration, removing adapter contamination, reads with over 10% uncertain nucleotides, and reads where low-quality bases exceeded 50% of the sequence. Post-filtration, the data were aligned to the mm10 mouse genome using the STAR software. For gene expression analysis, transcripts were quantified using the FPKM (Fragments Per Kilobase of transcript per Million mapped reads) metric.

To prepare for principal component analysis (PCA) of the time course data, the FPKM table was processed with the scanpy Python library. This involved filtering genes present in less than one cell (sc.pp.filter_genes(adata, min_cells=1)) and transforming the data via natural logarithm (sc.pp.log1p(adata)). PCA was then conducted to identify key components of variation, extracting and sorting the top loadings for each principal component based on both absolute and actual values. The final gene loadings and gene names were saved in CSV format, allowing for detailed downstream analysis. The top most highly loaded 400 genes from PC1 were then subjected to GO analysis using g:Profiler (https://biit.cs.ut.ee/gprofiler/gost).

To analyze the enrichment of differential expressed genes in wildtype compared to *Hprt* KO RAW 264.7 cells, Novogene performed differential expression analysis of the two conditions by using the DESeq2 R package, with the significance criteria of padj<0.05 and log2FoldChange|>1. The gene lists of significance were uploaded to DAVID and only GO terms at the most specific resolution (GO-BP,MF,CC at level 5) were considered for annotation. The Benjamini metric (adjusted p-val for controlling false discovery rate) for the top 10 terms was projected into negative log10 space using numpy for the log transformation and matplotlib was used for the visualization.

### Isotopic labeling

For stable isotope labeling with glutamine or glucose, cells were incubated with media containing the designated tracer γ-^15^N-glutamine (Cambridge Isotope Laboratories: NLM-557-1), or U-^13^C-glucose (Cambridge Isotope Laboratories: CLM-1396-5), which replaces the unlabeled glutamine or glucose in regular media at the same concentration. For stable isotope labeling with supplemented labeled inosine or formate, 50 µM U-^15^N-inosine (Cambridge Isotope Laboratories: NLM-4264-0.05) or 1mM ^13^C formate (Sigma-Aldrich: 279412) was added to the regular media. Cells were cultured with labeled media for the duration indicated in figure legend. Data from labeling experiments was adjusted for natural abundance of ^15^N and ^13^C.

### Immunoblotting

For protein collection, cells were washed twice with DPBS and lysed using RIPA buffer containing phosphatase and protease inhibitors (Thermo Fisher Scientific: A32957; A32953). Whole cell lysate was passed through a 27G needle (BD Biosciences: 305109) and centrifuged at 12000xg at 4°C. Protein content in the supernatant was quantified using a BCA assay kit (Thermo Fisher Scientific: 23223). Protein samples were heat denatured and separated on 4–12% Bolt Bis-Tris gels (Thermo Fisher Scientific: NW04125BOX). The protein was then transferred onto a 0.2μM nitrocellulose membrane (VWR: 27376-991). Membranes were blocked in 5% nonfat dairy milk in TBS-T, and probed for the following primary antibodies overnight:

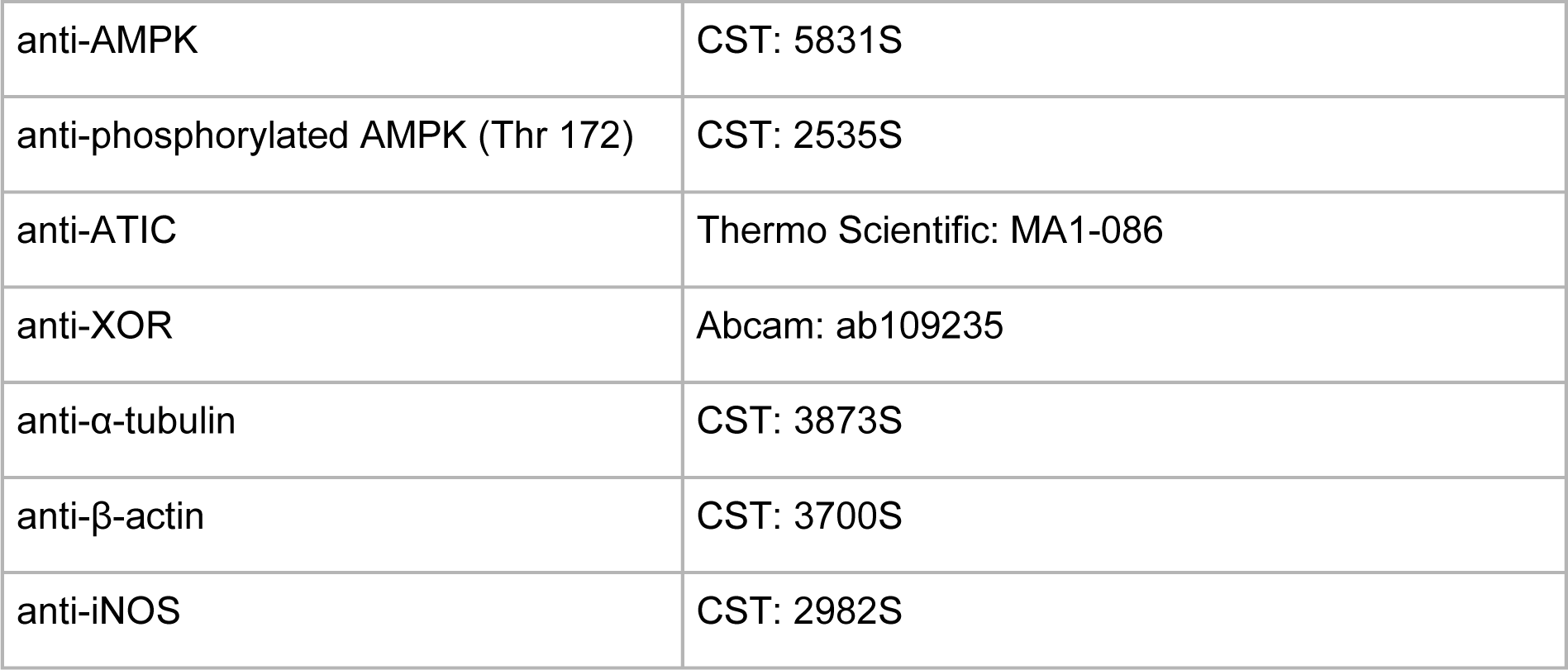

The following day, membranes were washed with TBS-T and then probed with Li-Cor secondary antibodies (goat anti-rabbit 800CW, goat anti-mouse 680RD [Li-Cor Biosciences: 926-32211; 926-68070]). Membranes were imaged with a Li-Cor Odyssey ClX and quantified using ImageStudio Lite software.

### Generation of knock out cell lines

Genetic knockouts were generated using the Alt-R RNP system in low passage wildtype RAW 264.7 cells following the manufacturer’s instructions. The system consists of Alt-R S.p. HiFi Cas9 Nuclease V3 (IDT: 1081060), Alt-R CRISPR–Cas9 gRNA (IDT), and ATTO 550 (IDT: 1075927).

The following gRNA sequences were used:

XOR: GAAGACGTTGCGTTTTGAAG

HPRT: ACGGGGGACATAAAAGTTAT

*Nos2*^-/-^ RAW 264.7 cells that were used were previously generated as described (Seim et al., 2023).

After the RNP complexes were formed, cells were transfected by electroporation using the Amaxa Nucleofector system Kit V (Lonza: VCA-1003). The following day, cells positive for fluorescent tracrRNA were single-cell sorted into individual wells of a 96-well plate containing RPMI, 25mM HEPES, 2mM glutamine, 20% fetal bovine serum. Single-cell colonies were subsequently expanded. To validate gene knockout from single-cell sorted colonies, genomic DNA was extracted using the QIAamp DNA Mini Kit (Qiagen: 51304) and a region around the CRISPR cut site target was amplified using the following primers containing M13 tags for sequencing.

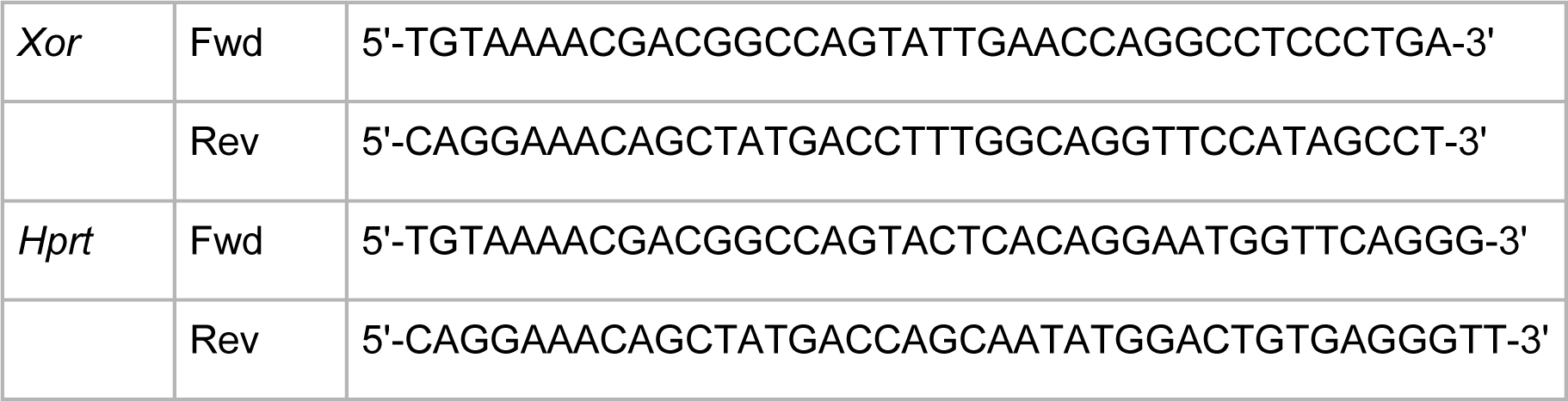

Knockouts were verified by Sanger sequencing (Genewiz) of the PCR product after column purification (Thermo Fisher Scientific: K0701).

### Oxygen consumption rate measurement

Oxygen consumption rates (OCR) were measured using an Agilent XF Seahorse Extracellular Flux Analyzer according to the manufacturer’s instructions. Cells were plated at a seeding density of 6×10^4^ cells per well in normal cell culture media. Before measuring OCR, cells were switched over to XF DMEM Medium, pH 7.4 (Agilent: 103193-100) containing 11.11mM glucose, 1% antibiotics, 5mM HEPES, 2mM glutamine, and 20ng/mL mCSF, without phenol red, sodium bicarbonate, or sodium pyruvate. Data was blanked from wells containing only media.

### Cell migration

Cell migration across transwells was performed as previously described (Chen et al., 2018) using an Incucyte live cell imager. For migration assays, Matrigel (Fisher Scientific: CB-40234) was diluted to 50ug/mL in DPBS to coat the top and bottom sides of the inserts of an Incucyte Clearview 96-well plate (Sartorius: 4582). After 1h, residual Matrigel solution was aspirated and RAW 264.7 cells (wildtype and *Hprt* KOs) were plated in RPMI containing 2mM glutamine and 0.5% dialyzed fetal bovine serum (assay medium) at a seeding density of 9,000 cells per well on the top chamber. The cells were allowed to settle and adhere to the top membrane for 1h. Afterwards, the chemoattractants LPS or C5a (R&D Systems: CB-40234) were added to assay medium in the bottom reservoir at concentrations of 1000ng/mL or 200ng/mL, respectively. Cell migration was imaged continually for 48h. Data analysis was performed using the Incucyte Chemotaxis module that quantified the number of cells present on the bottom side of the transwell at each timepoint imaged.

### Pyroptosis assay

For pyroptosis assays, 30,000 RAW 264.7 cells (wildtype and *Hprt* KOs) per well were plated in 96-well plates, in either regular culture media (un-primed control) or media containing LPS+IFNγ and incubated for 24h. Three hours before induction of pyroptosis, the media was refreshed with the addition of 250nM Cytotox green (Sartorius). Pyroptosis was induced with 10μM nigericin (Cayman Chemical: 11437) and the cells were subsequently imaged over a timecourse using an Incucyte S3 Live-Cell imager (Sartorius). Pyroptosis was measured based on a cell-by-cell classification analysis that quantified the percentage of green positive cells out of the total number of cells imaged.

### Analysis of Toxoplasma gondii growth

*T. gondii* tachyzoites were cultured in human foreskin fibroblasts in Dulbecco’s modified Eagle’s medium (Gibco: 11960-051) supplemented with 10% fetal bovine serum, 2mM glutamine, and 1% antibiotics at 37°C with 5% CO_2_. *T. gondii* growth in macrophages was measured by two imaging-based methods. To determine overall *T. gondii* burden in live culture over time, 5×10^4^ viable RAW 264.7 macrophages (wildtype, *Xor* KO or *Hprt* KO) were seeded in each well of a 48-well plate and infected with 5×10^4^ mCherry ME49 (type II) tachyzoites. The plate was incubated at 37°C and 5% CO_2_ and imaged using an Incucyte Live-Cell imager (Sartorius). The red channel used an acquisition time of 400 milliseconds for all replicates and samples. The parasites (red) were measured at a threshold of 0.90 Red Calculated Units with surface-fit segmentation. To image *T. gondii* proliferation at higher resolution and analyze parasite number in vacuoles by immunofluorescence, a total of 1×10^5^ viable macrophages were seeded on glass coverslips and infected with 1×10^5^ mCherry ME49 (type II) tachyzoites. At 24h post infection, cells were fixed with 4% paraformaldehyde, permeabilized with 0.5% Triton X-100 (Fisher Scientific: BP151-500) and blocked with 5% BSA. Cells were incubated with primary antibody against *T. gondii* surface antigen 1 (SAG1, rabbit, 1:300; Genetex GTX38936) overnight at 4°C and then thoroughly washed with DPBS. Secondary antibody against Alexa Fluor 594 (goat anti-rabbit, 1:1000; Thermo Scientific: A11012) was incubated for 2h at room temperature. Cells were counterstained with DAPI (Sigma D9564) and mounted using VECTASHIELD antifade Mounting Medium (Vector Laboratories: H-1000). Cells were imaged using an epifluorescence microscope (Imager.M2, Carl Zeiss, DEU). Vacuoles, parasites per vacuole, infected macrophages, and non-infected macrophages were counted in 10 random fields per coverslip. Each timepoint was performed on triplicate coverslips per condition. All coverslips were blinded before counting.

## Acknowledgements

This work is supported by the Morgridge Institute for Research, NIH grant R35 GM147014 (J.F.), R01 AI172874 (L.J.K.), and R56 AI158958 (J.F.). G.L.S. was supported by NRSA Individual Predoctoral Fellowship F31AI152280. B. J. E-F. was supported by training grant T32 AI007414. Flow cytometry and single cell sorting were performed with the instrument and assistance of UWCCC Flow Lab Core, supported by Cancer Center Support Grant P30 CA014520. The authors thank Matt Stefely for assistance in figure editing and Alicia Williams for text editing.

